# GRK specificity and Gβγ dependency determines a GPCR’s potential in biased agonism

**DOI:** 10.1101/2023.07.14.548990

**Authors:** Edda S. F. Matthees, Jenny C. Filor, Natasha Jaiswal, Mona Reichel, Noureldine Youssef, Julia Drube, Amod Godbole, Carsten Hoffmann

## Abstract

G protein-coupled receptors (GPCRs) are mainly regulated by GPCR kinase (GRK) phosphorylation and subsequent β-arrestin recruitment. Recently, it was shown that GPCRs differentially depend on GRK2/3, GRK2/3/5/6 or GRK5/6 for their regulation. The four ubiquitously expressed GRKs are classified into the cytosolic GRK2/3 and the membrane-tethered GRK5/6 subfamily. In vitro studies revealed that GRK2/3 interact with the membrane-localized G protein βγ-subunits. Yet, the role of this interaction as crosslink between G protein activation and β-arrestin binding to GPCRs remained strongly underappreciated. Here we systematically show that the Gβγ–GRK2/3 interaction is key for these GRKs to mediate β-arrestin2 binding to Gs-, Gi- and Gq-coupled GPCRs. In our GRK2/3/5/6 knockout cells, without endogenous GRK background, the utilized GRK2/3 mutants devoid of the Gβγ interaction site significantly diminished β-arrestin2 recruitment to the beta-2 adrenergic receptor (b2AR), muscarinic M2 and M5 acetylcholine receptors (M2R, M5R). This effect was overwritten by artificially tethering GRK2/3 via a CAAX motif to the plasma membrane independently of free Gβγ. Hence, the membrane recruitment is crucial for GRK2/3-mediated β-arrestin2 binding to GPCRs, which is naturally induced via the Gβγ interaction. This connects the β-arrestin interaction for GRK2/3-regulated receptors inseparably with the associated G protein activation. We outline a theoretical framework of how GRK dependence on free Gβγ can determine a GPCR’s potential in biased agonism. Due to this inherent cellular mechanism for GRK2/3 recruitment and receptor phosphorylation, we propose that it will likely be mechanistically unattainable to create β-arrestin-biased ligands for the subgroup of GRK2/3-regulated GPCRs, while GRK5/6-mediated receptor regulation is independent from Gβγ availability. Accordingly, one should first determine the GRK specificity of a GPCR to ultimately assess the receptor’s potential for the development of biased ligands.

## Introduction

Upon agonist stimulation, a G protein-coupled receptor (GPCR) induces structural changes in the heterotrimeric G protein to induce G protein-dependent signalling^1^. GPCR kinases (GRKs) play a critical role in regulating this signalling, as they are the major class of kinases to phosphorylate active GPCRs to initiate receptor desensitization by enhancing arrestin recruitment^2-4^. Due to initial reports that unwanted side effects of opioid treatment were reduced in β-arrestin2 knockout mice^5^, multiple studies aimed to introduce bias between G protein- or β-arrestin-mediated GPCR pathways^6-8^. In general, biased signaling describes the capability of a ligand at a given receptor to preferentially trigger one signaling pathway over another when compared to a reference ligand^9^. In a ternary complex of a ligand, a receptor and a transducer, theoretically any of the three components can be the cause of an observed signaling bias^10^. Currently, the mechanistic aspects of biased agonism are described from two perspectives. The first aspect would be the structural component, which is inherent to possible conformations of a GPCR, coupling differentially to distinct cellular signaling pathways, and can be controlled ideally by selective ligands^11,12^. The second component contains the respective transducer and is often unknown and encoded with the factor τ, which is used to express biased factors for ligands and contains the cellular contributors of biased signaling^13^. However, which overarching mechanism for GPCRs generally determines the shift in balance between G protein and β-arrestin pathways remains unclear.

With the aim of unravelling one such component in this mechanism, we utilized our recently published CRISPR/Cas9-edited HEK293 cells devoid of GRK2/3/5/6 (ΔQ-GRK cells)^14^ to observe that GPCRs show GRK selectivity to mediate β-arrestin recruitment. Of particular importance was the finding that some receptors relied solely on GRK2/3-mediated regulation, whereas others did not show any preference among the four ubiquitously expressed GRKs (GRK2/3/5/6-regulated) (Tab. 1)^14^. Recent studies additionally indicated the existence of a third category of only GRK5/6-dependent receptors (Tab.1)^15,16^.

**Tab.1:** Overview of classified GRK2/3-, GRK2/3/5/6-, GRK5/6-regulated GPCRs in current literature.

| GRK2/3 | GRK2/3/5/6 | GRK5/6 |
| --- | --- | --- |
| M2R<sup>14</sup> | AT1R<sup>14,17</sup> | C5aR2<sup>15</sup> |
| M4R<sup>14</sup> | C5aR1<sup>14,15</sup> | ACKR3<sup>16</sup> (Youssef et al., in preparation) |
| M5R<sup>14</sup> | CCR2<sup>15</sup> |  |
| MOP<sup>14,18</sup> | M3R<sup>14</sup> |  |
|  | PTH1R<sup>14</sup> |  |
|  | V2R<sup>14</sup> |  |
|  | b2AR<sup>14</sup> |  |

This intriguing GRK selectivity led us to revisit the basic molecular mechanisms of how GRKs are stabilized at the plasma membrane to interact with GPCRs and to evaluate this as a crucial and central molecular mechanism of biased signalling. As previously reported, the cytosolic GRK2/3 rely on free Gβγ subunits to translocate to the membrane at the site of active receptor^19-22^. Only after this Gβγ-mediated GRK2/3 recruitment, can the receptor be phosphorylated to enhance β-arrestin2 recruitment^23^. Conversely, the membrane localization of GRK5/6 is independent of free Gβγ proteins^24-26^. Utilizing our ΔQ-GRK cells, we systematically evaluated that indeed Gβγ interaction is crucial for GRK2/3-mediated β-arrestin recruitment to Gs-, Gi- and Gq-coupled GPCRs whereas interaction with Gα is negligible.

Our findings could have significant impact on studies of biased agonism for G protein *versus* β-arrestin pathways since our data imply that all GRK2/3-mediated β-arrestin effects are indeed G protein-(βγ)-dependent and only GRK5/6-mediated effects on β-arrestin are G protein-independent. In this study, we outline a theoretical framework of how GRK dependence on free Gβγ can determine a GPCR’s potential in biased agonism.

## Results

### GRK2-Gβγ binding is essential to recruit β-arrestin2 to b2AR

To investigate the G protein dependency of GRK2-mediated β-arrestin2 recruitment to activated GPCRs, we used a variety of GRK2-mutants with low affinity binding towards the G protein subunits and our quadruple knockout cell line devoid of ubiquitously expressed GRK isoforms 2/3/5 and 6 (ΔQ-GRK cells)^14^. As depicted in Fig. 1A, these mutant versions disrupt the interaction of the GRK2 with Gαq (D110A)^20^, the interaction with Gβγ (R587Q)^27,28^ or both (double mutation at D110A and R587Q)^23^. We also created CAAX-tagged versions of GRK2 and the above mutants^29^ to localize the cytosolic GRKs permanently to the plasma membrane and thus overwrite the requirement on binding to active G protein subunits.

**Fig.1:**
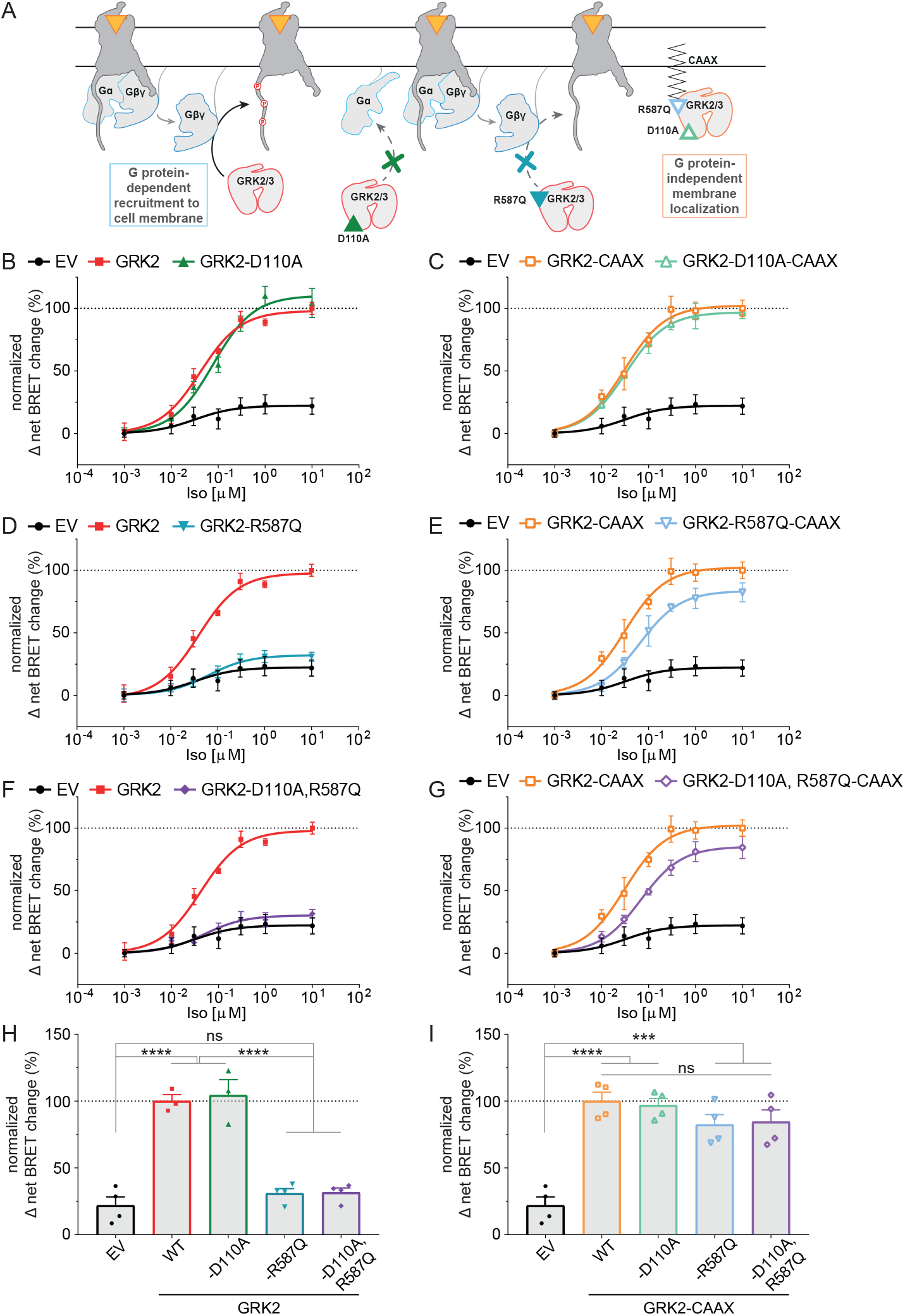
GRK2-mediated β-arrestin2 recruitment to the beta-2 adrenergic receptor (b2AR) is dependent on the membrane localization of the GRK. **A**, Schematic representation of the utilized GRK mutants D110A (interrupting the GRK2/3–Gα interaction), R587Q (interrupting the GRK2/3– Gβγ interaction) and the double mutant (D110A, R587Q), also as versions with a CAAX box to localize GRK2/3 to the plasma membrane independent of the G protein interaction. **B-G**, Isoprenaline (Iso)-induced Halo-Tag-β-arrestin2 recruitment to b2AR-NanoLuciferase (NLuc) in GRK2/3/5/6-depleted quadruple knockout HEK293 (ΔQ-GRK) cells in absence of the ubiquitously expressed GRKs (empty vector (EV)-transfected) and in presence of wild type GRK2 or either GRK2-D110A (B), GRK2-R587Q (D) or GRK2-D110A,R587Q (F). The same experiment was performed with the corresponding GRK2-CAAX versions (C, E, G). All data are shown as Δ net BRET change in percent of at least *n=*3 independent experiments ± SEM, normalized to the maximum response with GRK2 (B, D, F) or GRK2-CAAX (C, E, G). The curves in absence of ubiquitously expressed GRKs (EV), GRK2 and GRK2-CAAX are shown multiple times to allow direct comparisons. **H, I**, Normalized BRET data of the highest stimulation of B-G are displayed as bar graphs and statistical differences were tested using one-way ANOVA, followed by a Turkey’s test (ns not significant; * p < 0.05; ** p < 0.01; *** p < 0.001; **** p < 0.0001). Detailed statistical results are provided in Suppl. Tab. 1.

First, we investigated the dependency of β-arrestin2 on interaction between activated G proteins and GRKs when recruited to the prototypical Gs-coupled beta-2 adrenergic receptor (b2AR). Previously, we demonstrated that each GRK2/3/5 and 6 can enhance recruitment of β-arrestin to b2AR upon agonist stimulation^14^. Here, we re-introduced the various GRK2 mutants mentioned above in combination with Nanoluciferase (NLuc)-tagged b2AR and Halo-tagged β-arrestin2 in our ΔQ-GRK cells and monitored b2AR–β-arrestin2 interaction upon stimulation with the agonist isoproterenol (Iso).

For data visualization, we plotted the normalized net BRET change as concentration-response curves (Fig. 1B-G) as well as in bar graphs for the highest agonist concentration to compare the effect of the different GK2 versions (Fig. 1H-I, details on statistical analysis are provided in Suppl. Tab. 1). The GRK2-D110A mutant has been described to disrupt binding of GRK2 specifically with Gαq^20,22^. As anticipated our findings show that this mutant did not negatively influence β-arrestin2 recruitment to the Gs-coupled b2AR as compared to wild type (WT) GRK2 (Fig. 1B and 1H). The introduction of the CAAX motif to the WT GRK2 or the GRK2-D110A did not affect its ability to mediate β-arrestin2 recruitment to the b2AR (Fig. 1C and 1I, Suppl. Fig. 1, Suppl. Tab. 2). These findings imply that the interaction between GRK2 and the Gα subunit is secondary. Interestingly, disruption of GRK2 binding to free Gβγ subunits using the GRK2-R587Q mutant did significantly reduce β-arrestin2 recruitment to amplitudes seen in the condition without endogenous GRK2/3/5/6 expression (Figure 1D and 1H). The localization of this GRK2-R587Q mutant on the plasma membrane restored β-arrestin2 recruitment to similar amplitude as seen for the WT GRK2-CAAX (Figure 1E and 1I) further highlighting that removing the dependence of GRK2 on its interaction with activated Gβγ subunits can circumvent receptor phosphorylation and hence β-arrestin recruitment. In line with these findings, the double mutant (GRK2-D110A,R587Q) also drastically reduced β-arrestin2 recruitment compared to WT GRK2 (Figure 1F and 1H) whereas removing the dependence on activated Gβγ subunits by using plasma-membrane localized GRK2 double mutant restored β-arrestin2 recruitment as seen for the WT GRK2-CAAX (Figure 1G and 1I).

With this observation that GRK2-regulated β-arrestin2 recruitment to a Gs-coupled receptor would always be preceded by free Gβγ subunits, we investigated whether this is independent of the Gβγ source.

### GRK2-Gβγ interaction is necessary to mediate β-arrestin2 recruitment to the muscarinic M2R and M5R

To this aim, we then tested the prototypical Gi-coupled muscarinic M2 receptor (M2R), which has been shown by us to be dependent on only GRK2/3 to mediate β-arrestin recruitment^14^. We tested the dependency of β-arrestin2 recruitment to this receptor on GRK–G protein interactions by using similar approaches and mutants as described above (Fig. 2A and B). As seen for the b2AR (Fig. 1), the disruption of the binding of GRK2 to free Gβγ subunits by using the GRK2-R587Q mutant significantly reduced β-arrestin2 recruitment to the M2R, an effect which was shared also with the double mutant (Figure 2A). Targeting GRK2 to the membrane using the CAAX-tag led to a reduced dynamic change of β-arrestin2 recruitment to this receptor (Suppl. Fig. 2, Suppl. Tab. 5). Nevertheless, utilizing the GRK2-CAAX version allowed the M2R to recruit β-arrestin2 by surpassing the need of free Gβγ subunits as shown by the GRK2-R587Q-CAAX mutant (Figure 2B).

**Fig.2:**
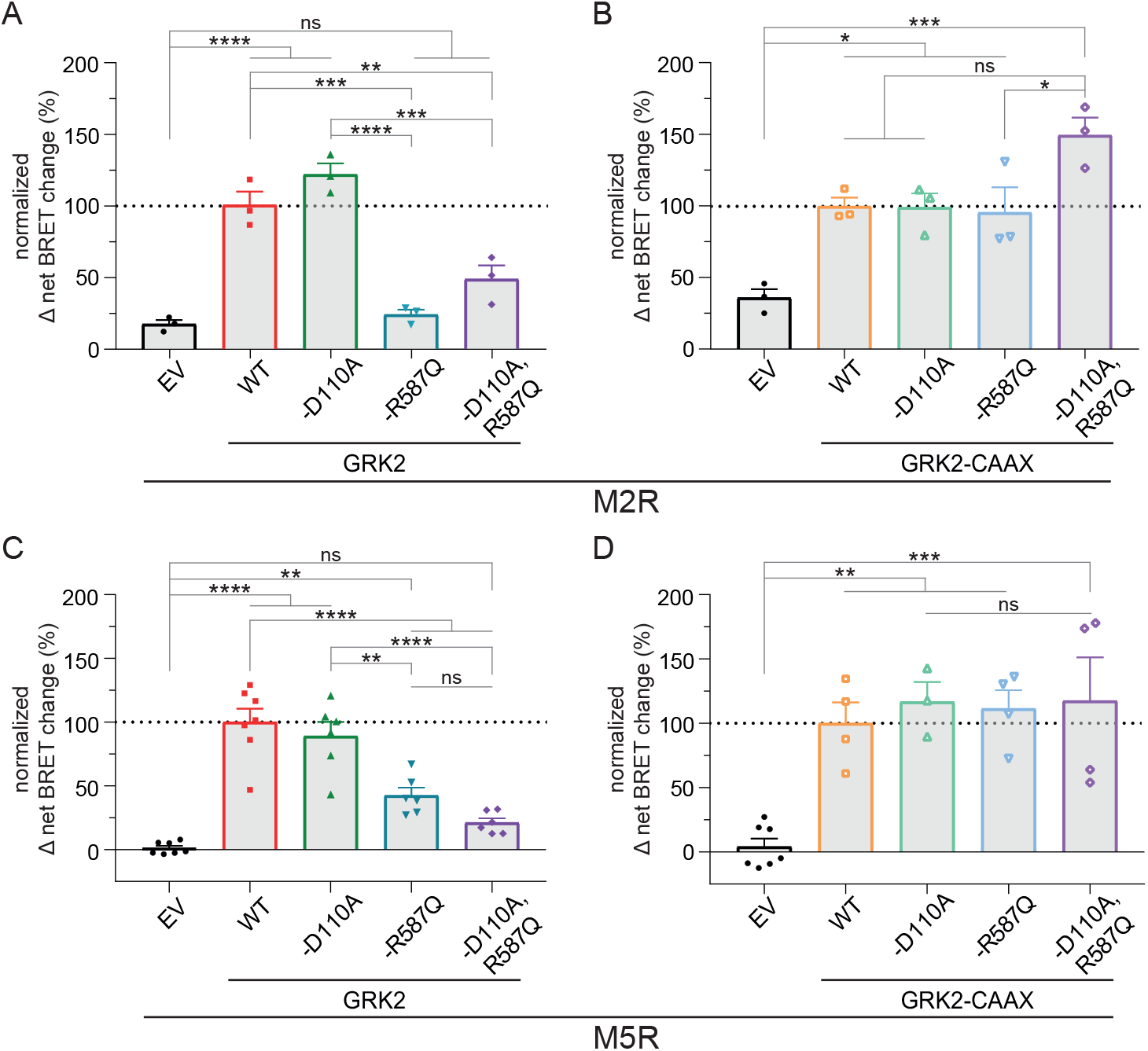
β-arrestin2 recruitment to the muscarinic M2 and M5 acetylcholine receptors (M2R, M5R) is dependent on the ability of the GRK to be recruited to the membrane. **A-D**, Halo-Tag-β-arrestin2 recruitment to M2R-NLuc (A, B) or M5R-NLuc (C, D) was measured in ΔQ-GRK cells in absence of GRKs (EV-transfected) and in presence of wild type GRK2, GRK2-D110A, GRK2-R587Q, GRK2-D110A,R587Q or their respective membrane-tethered versions via a CAAX box (B, D). Normalized Δ net BRET change (%) upon stimulation with 100 μM of Acetylcholine (ACh) is shown of at least *n=*3 independent experiments ± SEM, normalized to GRK2 (A, C) or GRK2-CAAX (B, D). The data measured in absence of GRKs (EV) is shown in both graphs each (A, B for M2R; C, D for M5R) for direct comparison. Statistical differences were tested using one-way ANOVA, followed by a Tukey’s test (ns not significant; * p < 0.05; ** p < 0.01; *** p < 0.001; **** p < 0.0001). Detailed statistical results are provided in Suppl. Tab. 3 (M2R) and Suppl. Tab. 4 (M5R).

To test the dependency of β-arrestin2 on GRK–G protein interaction for a Gq-coupled receptor, we investigated the muscarinic M5 receptor (M5R), which we have shown to be GRK2/3-regulated with respect to β-arrestin recruitment^14^. As seen for the previous receptors, disruption of GRK2 binding to active Gβγ subunits significantly reduced β-arrestin2 recruitment (Fig. 2C). Interestingly, disruption of GRK2 binding to the Gαq subunit did not reduce β-arrestin2 recruitment (Fig. 2C). That is in line with previous reports that GRK2-Gαq interactions is secondary to GRK-Gβγ interaction^29^. These findings now performed without an endogenous GRK background clearly unravel the role of free Gβγ subunits on the ability of GRK2 to regulate receptors. As seen for the other receptors, surpassing the need of free Gβγ subunits by using CAAX-tagged GRK2, the recruitment of β-arrestin2 to M5R was restored to the same levels of WT GRK2-CAAX (Fig. 2D). As seen for the M2R, introducing the CAAX motif to the GRK2 reduced the agonist-dependent dynamic change of mediated β-arrestin2 recruitment comparted to WT GRK2 (Suppl. Fig. 3, Suppl. Tab. 6), indicating that the functional receptor interaction might be different for the membrane-tethered GRK2 with these GPCRs as opposed to the b2AR (Suppl. Fig. 1). Nevertheless, the relative observation held true for all the compared receptors.

In summary, the data with Gs-, Gi- and Gq-coupled receptors point out a general mechanism in which free Gβγ subunits play a critical role in translocating cytosolic GRK2 to the plasma membrane. This Gβγ dependency was generally circumvented by introducing the membrane localization motif (CAAX) to the GRK2.

### GRK3 is similarly dependent on membrane-recruitment to phosphorylate GPCRs as family member GRK2

Next, we tested the dependency of β-arrestin2 on GRK3–G protein interactions since GRK3 is the other subfamily member and yet less investigated kinase in the field of receptor regulation. We obtained similar results with GRK3 and its Gαq-(D110A) or Gβγ-(R587Q) interaction mutants, as well as the double mutant (Figure 3 A, C, E). The introduction of the CAAX motif reduced the ability of the GRK to mediate β-arrestin2 recruitment to these receptors, similar to what we observed for GRK2 (Suppl. Fig. 4, Suppl. Tab. 10). In contrast to GRK2, the GRK3-D110A,R587Q-CAAX mutant version generally interfered with the extend of mediated β-arrestin2 recruitment stronger compared to the other GRK3-CAAX versions (Fig. 3 B, D, F).

**Fig.3:**
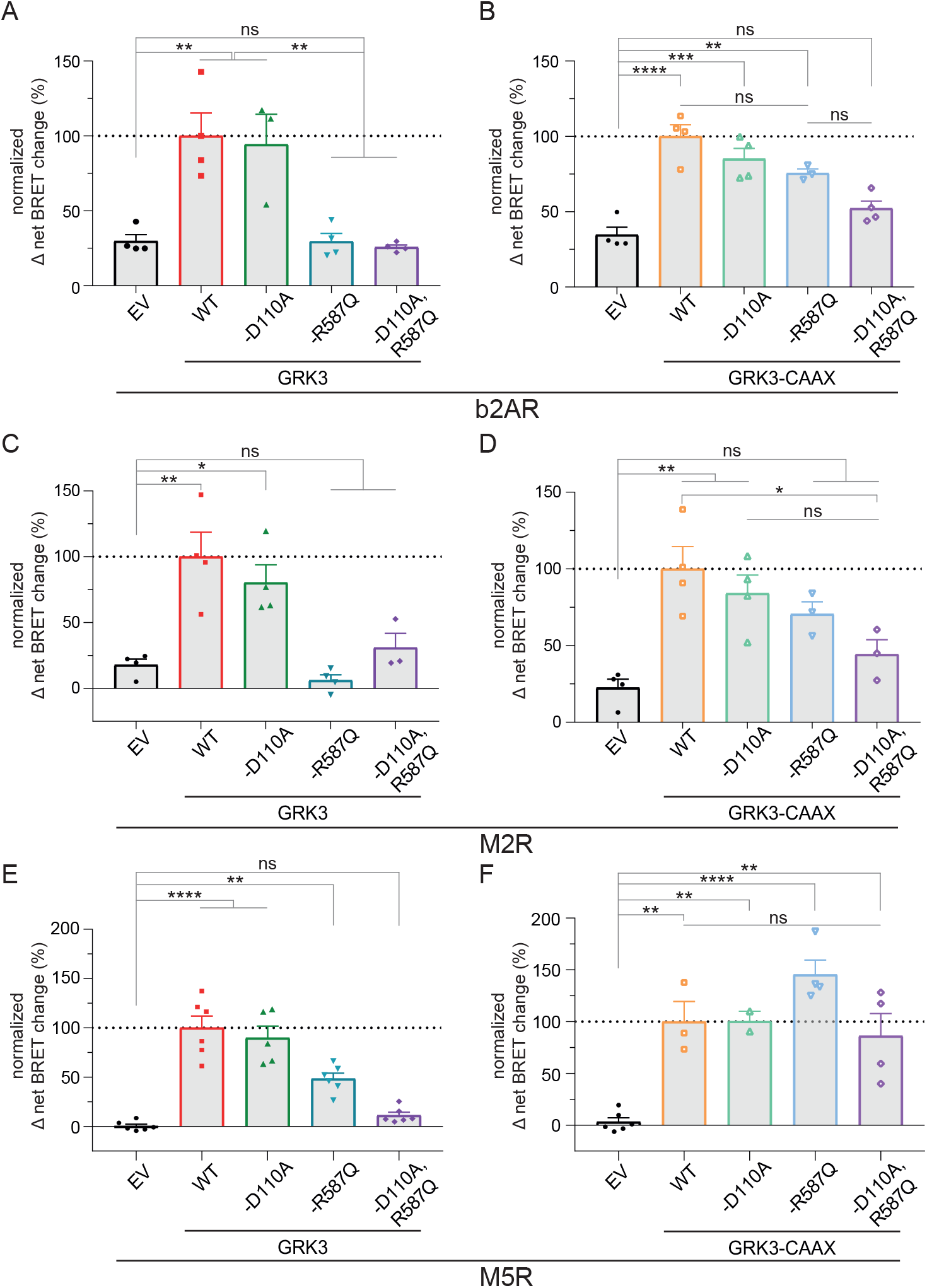
GRK3-mediated β-arrestin2 recruitment to b2AR, M2R and M5R displayed a similar dependency as GRK2 on the ability of the GRK to be recruited to the membrane. **A-F**, Halo-Tag-β-arrestin2 recruitment to b2AR-Nluc (A, B), M2R-NLuc (C, D) or M5R-NLuc (E, F) was measured in ΔQ-GRK cells analogous to GRK2 in absence of GRKs (EV-transfected) and in presence of wild type GRK3, GRK3-D110A, GRK3-R587Q, GRK3-D110A,R587Q or their respective membrane-tethered versions via a CAAX box (B, D, F). Normalized Δ net BRET change (%) upon stimulation with 10 μM Iso (A, B) or 100 μM of ACh (C-F) is shown of at least *n=*3 independent experiments ± SEM, normalized to GRK3 (A, C, E) or GRK3-CAAX (B, D, F). The data measured in absence of GRKs (EV) is shown in both graphs respectively (A, B for b2AR; C, D for M2R; E, F for M5R) for direct comparison. Statistical differences were tested using one-way ANOVA, followed by a Tukey’s test (ns not significant; * p < 0.05; ** p < 0.01; *** p < 0.001; **** p < 0.0001). Detailed statistical results are provided in Suppl. Tab. 7 (b2AR), Suppl. Tab. 8 (M2R) and Suppl. Tab. 9 (M5R).

This potentially indicates some differences between the two closely related GRK isoforms. Still, the findings for the cytosolic GRK3–Gβγ interaction mutants strongly indicate how receptors which are solely regulated by GRK2/3 are largely dependent on agonist-dependent G protein rearrangement.

### GRK2/3-mediated β-arrestin2 recruitment is diminished by competitive inhibition of GRK2/3-Gβγ-interaction

In this study, we show that GRK2/3-mediated β-arrestin2 recruitment is dependent on the interaction of GRK2/3 with Gβγ by mutation of the Gβγ interaction site at GRK2/3 (Fig. 1-3). To investigate this principle using a different approach, we utilized the C-terminal domain of GRK2, referred to as bARK-CT, as a described competitive inhibitor for GRK2/3–Gβγ interaction^27,30^. The identified GRK2–Gβγ interaction site R587^27,28^ is included in the bARK-CT peptide (Fig. 4 A). We investigated the influence of bARK-CT co-expression on GRK2/3-mediated β-arrestin2 recruitment to the M5R (Fig. 4B). Of the investigated receptors, the M5R was chosen since it is described as a GRK2/3-regulated GPCR, also under endogenous expression levels of the ubiquitously expressed GRKs unlike the M2R^14^. The b2AR was omitted as it is regulated by GRK2/3/5/6^14^. Based on this, the experiment was performed at endogenous expression levels of GRKs in our HEK293 control cells. To test the influence of bARK-CT on GRK2/3-mediated β-arrestin2 recruitment 0.5 μg or 1 μg of bARK-CT was transfected as indicated in addition to NLuc-tagged M5R and Halo-tagged β-arrestin2, as described above.

**Fig.4:**
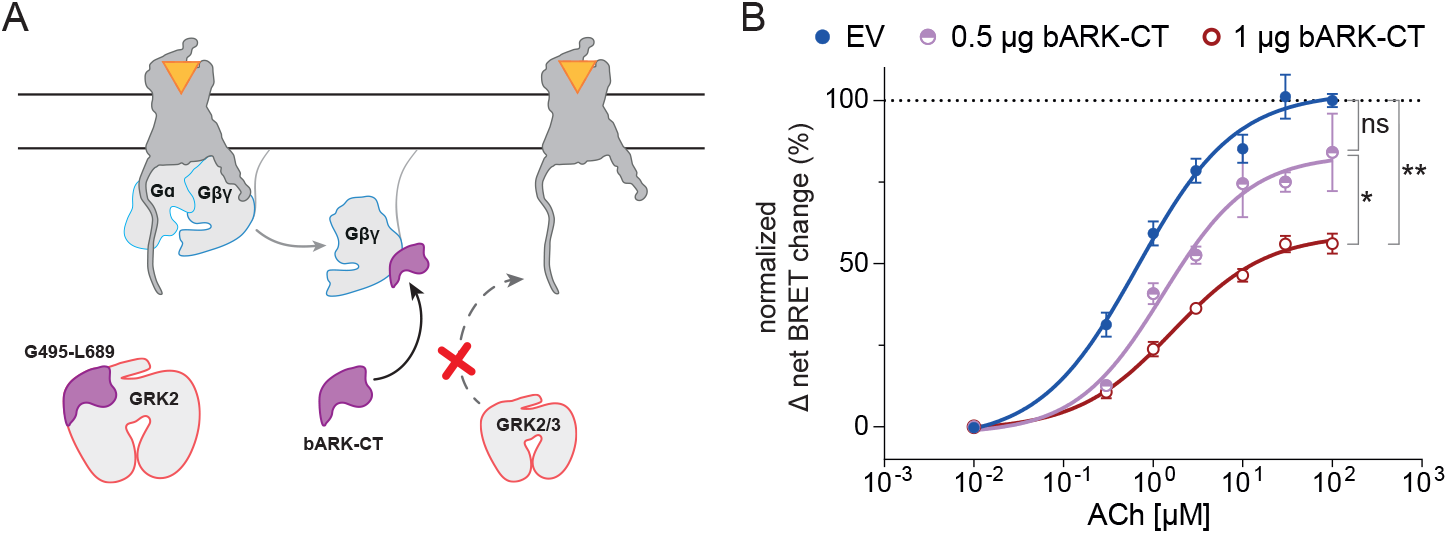
bARK-CT reduces β-arrestin2 recruitment to the muscarinic M5 acetylcholine receptor (M5R). **A**, Schematic representation of the bARK-CT-mediated mechanism as an inhibitor of GRK2/3 recruitment to the M5R. The bARK-CT fragment of GRK2 includes the GRK2-Gβγ interaction site (R587) and therefore competes with GRK2/3 for the binding. **B**, Halo-Tag-β-arrestin2 recruitment to M5R-NLuc was measured in CRISPR/Cas9 Control cells, expressing all GRKs at endogenous levels, in absence (empty vector (EV)-transfected) or presence of different co-transfected amounts of bARK-CT (as indicated). Normalized Δ net BRET change (%) upon stimulation with the indicated concentrations of Acetylcholine (ACh) is shown of at least *n=*3 independent experiments ± SEM, normalized to EV. Statistical differences were tested using one-way ANOVA, followed by a Tukey’s test (ns not significant; * p < 0.05; ** p < 0.01). Detailed statistical results are provided in Suppl. Tab. 11.

Increasing amounts of transfected bARK-CT led to a gradual reduction of β-arrestin2 recruitment (Fig. 4B). This indicated that not just the mutation of the Gβγ interaction site in GRKs, but also the competitive inhibition of endogenous GRK2/3 by bARK-CT leads to a reduction of GRK2/3-mediated β-arrestin2 recruitment. These findings indicate that utilization of bARK-CT, which is a known inhibitor of Gβγ signaling, as a competitive GRK2/3 inhibitor would also affect β-arrestin2 recruitment.

## Discussion

In this study, we investigated the impact of the Gβγ interaction with GRK2/3 as a cellular mechanism for GRK2/3 translocation to and stabilization at the plasma membrane. We specifically studied the effect of this interaction on GPCR–β-arrestin complex formation. To this end, we utilized a number of previously described mutations in GRK2/3, which selectively interrupt either binding to Gβγ or Gαq or both and combined these mutations with a CAAX motif to tether the cytosolic GRKs to the plasma membrane independently of the disrupted Gβγ interaction. We performed these measurements in our recently published HEK293 knockout cell line devoid of endogenous GRK2/3/5/6 background^14^. Using this unique combination, we could show that the ability of GRK2 to mediate b2AR–β-arrestin interaction strongly depends on Gβγ binding (Fig. 1). Next, we demonstrated that this critical dependency of GRK2 on Gβγ binding is independent of the G protein coupling specificity of GPCRs as it is conserved for a Gs-, Gi- and Gq-coupled receptor (Fig. 1 and Fig. 2). This indicates that we did not observe specificity of the Gβγ source as seen for GIRK-channel activation via Gβγ of Gi proteins^31^ but rather a general impact of Gβγ. Analogously, we showed that this mechanism is also conserved for GRK3 at all investigated GPCRs (Fig. 3). In other words, GRK2/3 subfamily-mediated β-arrestin recruitment to GPCRs is severely determined by the ability of these GRKs to bind to free Gβγ subunits and hence, Gβγ availability. We additionally tested this observation in cells expressing the endogenous wild type GRK complement using bARK-CT as a known Gβγ inhibitor, which led to a dose-dependent inhibition of β-arrestin recruitment to the M5R (Fig. 4). Thus, unless other, yet unknown mechanisms of membrane recruitment substitute for this Gβγ interaction, G protein activation and GRK2/3-mediated β-arrestin recruitment are inseparably intertwined. When reviewing the literature, we found no prior evidence that this was comprehensively analyzed before in the absence of endogenous GRK background. However, it has been stated that a component of GRK2/3 recruitment requires GPCR signaling to liberate free Gβγ^32,33^. At the same time, it was noted that the potential impact of G protein activity on other signaling pathways regulated by arrestins yet needs to be determined but if confirmed, this would be hard to reconcile with the concept of G protein- or arrestin-biased ligands^34^.

Our findings presented in this article suggest that the GRK2/3 interaction with Gβγ is a fundamental mechanism and has far-reaching consequences for the global effort of creating biased agonists: we propose that the GRK isoform requirement of a GPCR to recruit β-arrestin determines the potential of creating biased agonists promoting β-arrestin recruitment without activating G proteins. Hence, we would like to broadly classify GPCRs based on their GRK-selectivity and connect this classification to a consequent possibility of creating biased agonists (Fig. 5).

**Fig.5:**
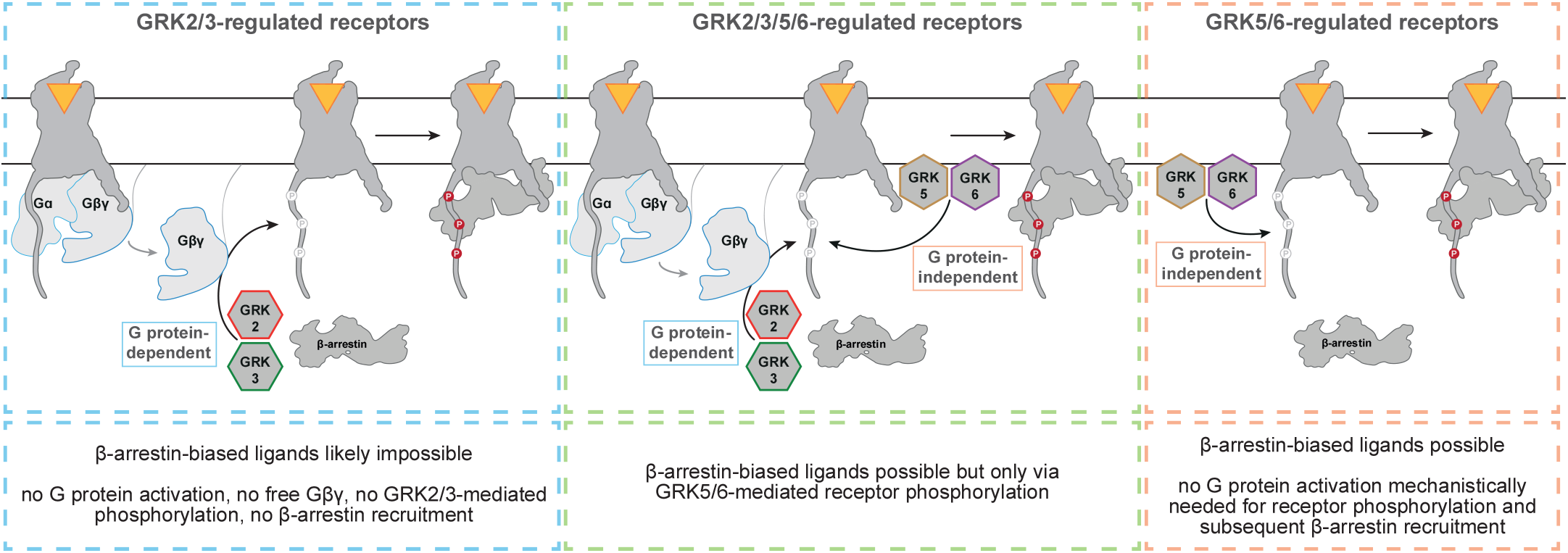
The GRK-dependency of each GPCR determines its potential in biased agonism. GPCRs can be grouped based on the GRKs involved in their regulation into GRK2/3-dependent, GRK5/6-dependent or GRK2/3/5/6-dependent receptors. As the membrane-localization of GRK2/3 is mediated via the interaction with Gβγ, the phosphorylation of the receptor and hence, the β-arrestin recruitment, are in fact G protein-dependent. Therefore, it will likely be mechanistically unattainable for this group of receptors to create β-arrestin-biased ligands that do not activate G proteins, because the phosphorylation by GRK2/3 is dependent on the availability of free Gβγ-subunits. This is not the case for GRK5/6-regulated receptors, as these GRKs are already membrane-tethered and not dependent on Gβγ for the recruitment to this receptor group. For receptors that are found to be GRK2/3/5/6-regulated, β-arrestin-biased ligands would convey their effects only via GRK5/6-induced receptor phosphorylation.

According to this GRK selectivity classification, we group GPCRs into GRK2/3-, GRK2/3/5/6- or GRK5/6-dependent receptors (Fig. 5). The creation of β-arrestin-biased agonists would be mechanistically unattainable for GRK2/3-regulated GPCRs, as the GRK2/3-mediated β-arrestin effects are indeed Gβγ-dependent. Partial agonists could lead to G protein-biased signaling if the subsequently available Gβγ is not sufficient to mediate efficient GPCR phosphorylation by GRK2/3. Further, receptors that are solely GRK5/6-regulated have intrinsically the highest possibility of obtaining β-arrestin-biased agonists since such agonists would not rely on the added necessity of a mechanistic G protein activation to induce phosphorylation and arrestin binding. Until now, the only two GRK5/6-regulated seven transmembrane receptors described in the literature are atypical ones, which are intrinsically β-arrestin-biased, naturally lacking G protein signaling^15,16^ (Youssef et al., in preparation). In other words, these receptors do not activate G proteins and therefore there would be no free Gβγ subunits available to mediate GRK2/3 recruitment to these receptors. Finally yet importantly, as β-arrestin recruitment to GRK2/3/5/6-regulated GPCRs is partly G protein-dependent (GRK2/3) and partly independent (GRK5/6), the creation of β-arrestin-biased agonists targeting GRK2/3/5/6-regulated GPCRs is generally possible. However, these would only mediate β-arrestin effects linked to GRK5/6 phosphorylation while GRK2/3 effects would be lacking since the absence of G protein activation would not deliver free Gβγ. Strong support for this hypothesis can be found in the literature. Two independent studies demonstrated clearly in GRK-knockout cell lines that the primarily Gq-coupled angiotensin-II (AngII) type 1 receptor (AT1R) is GRK2/3/5/6-regulated^14,17^. In Gq-knockout cells, this GRK specificity profile was shifted towards exclusive dependency on GRK5/6 for the balanced ligand AngII^17^. In this study, the same observation was made when wild type HEK293A cells were incubated with the chemical Gq inhibitor YM-254890. Using GRK family knockout cells, the authors also showed that the established β-arrestin-biased agonist TRV027 only mediates β-arrestin recruitment via GRK5/6 and no longer via GRK2/3 at the AT1R, as TRV027-mediated β-arrestin recruitment was only measurable in GRK2/3 knockout cells and not if GRK5/6 were knocked out^17^. We would argue that this originates from the lack of G protein activation and subsequent absence of free Gβγ to recruit GRK2/3 to the membrane. Hence, when designing β-arrestin-biased agonists one should keep in mind that the GRK5/6-mediated β-arrestin downstream effects might differ substantially from the functional outcome facilitated by GRK2/3/5/6-recruited β-arrestin^35,36^. Future studies should aim to further illuminate which β-arrestin-supported effects are carried out by GRK2/3 or GRK5/6 phosphorylation.

This G protein activation dependency of GRK2/3-mediated receptor regulation could be circumvented by alternative mechanisms of GRK2/3 recruitment to the membrane, demonstrated here by utilization of artificially membrane-tethered CAAX versions. In recent years, one study was published proposing to have observed such an alternative mechanism for GRK2/3 recruitment to the dopamine D2 receptor (D2R), independent of Gβγ interaction^37^. However, their strongest argument was the measurement of GRK2 recruitment in presence of the Gi inhibitor pertussis toxin (PTX). According to a comprehensive screen of GPCR–G protein selectivity published afterwards, the D2R was found to not only couple to Gi, but also to the less prominent Gz isoform^38^. This was recently confirmed using HEK293 cells with the knockout of six Gi isoforms including Gz. In this cell line, the D2R showed again strong coupling to Gz^39^. Since Gz is not sensitive to PTX^40,41^, this casts doubt on the full absence of free Gβγ when stimulating the D2R in presence of PTX. Rather than a Gβγ-independent mechanism, a different heterotrimeric G protein activated by the same receptor might have contributed to this finding. This raises the question whether utilized free Gβγ needs to originate from the same receptor. It has been shown for the atypical chemokine receptor ACKR3 that GRK2/3-mediated phosphorylation can be induced by overexpression of Gβγ and delivery of free Gβγ in the receptor vicinity via the hetero-dimerization with the G protein-activating CXCR4, if stimulated by CXCL12 as a shared agonist for both receptors^42^. In this case, the same ligand induced active conformations of the receptors, which might have enabled GRK2/3 to phosphorylate also the atypical ACKR3 even though the recruitment to the membrane was mediated via CXCR4.

Another layer of complexity is added to the translation of these findings into physiological contexts by possible variations of protein expression in cells. For example, a high combinatorial variability has been shown for more than 100 possible Gβγ dimer formations^43^ and GPCRs favorably interacting with specific Gβγ combinations^44^. Furthermore, the specific cellular function mediated by Gβγ also depends on the involved isoforms^45,46^. Additionally, it has been shown that Gγ subunits display distinct plasma membrane affinities, which influences different functional outcomes like the migration rate of macrophages^47^. The affinity to a receptor could differ for distinct combinations or in different tissues depending on the expression levels of the respective subunits. Future research can try to decipher whether or how specific Gβγ isoforms play distinct roles in the membrane recruitment of GRK2/3. Moreover, the differential tissue-specific expression of other involved proteins, like the GRKs themselves^35^, could influence biased signaling. Different GRK expression could lead to the possibility of an effector bias^48^ as only a subset of GRKs might be available to phosphorylate the receptor of interest, ultimately leading to different downstream consequences of GPCRs in specific tissues.

Our current findings in this conceptual work allowed this simple, previously impossible classification of GPCRs based on their GRK-selectivity. This classification of receptors into these three categories might be a key step in understanding if, which and how biased agonists will be possible. In summary, we propose an understanding of how β-arrestin-biased agonism works and how this biased agonism is highly dependent on which GRKs can regulate the activated receptor. To this aim, we pieced together all the available information on GRK2/3– Gβγ interactions and GRK2/3/5/6 cellular locations. We then used hitherto unavailable tools such as our recently published GRK knockout cell line in combination with established BRET assays to decipher the effect of Gβγ subunits on individual GRKs. Using these tools, we are putting forth a straightforward yet until now unobserved systematic understanding of the mechanism and the key players dictating biased agonism: the availability of free Gβγ subunits and the selectivity of receptors towards a specific set of GRKs. Ultimately, our findings hope to convey the simple message that we should consider whether there is a “GRK bias” of a receptor to assess its potential in G protein- and β-arrestin-biased signaling.

## Methods

### Cloning and construct origin

The transfected GRK constructs and receptors were of human and β-arrestin2 was of bovine origin. The NanoLuciferase (NLuc) and Halo-Tag genes were obtained from Promega and the plasmids expressing b2AR-, M2R- and M5R-NLuc and Halo-β-arrestin2 have been described before^14^. The bARK-CT construct was provided by Professor Silvio Gutkind. Plasmids expressing human full-length GRK2 and GRK3 in pcDNA3 have been described in Drube et al., 2022^14^ and plasmids expressing GRK2/3-D110A and GRK2/3-R587Q have been described in Jaiswal and Godbole et al., in preparation. The GRK2-D110A/R587Q double mutant was created by using GRK2-D110A as the template and 5’GGAGATCTTCGcCTCATACATCATGAAGGAGCTGCTGG as forward primer and 5’CCAGCAGCTCCTTCATGATGTATGAGGCGAAGATCTCC as reverse primer. GRK3 D110A/R587Q double mutant was created using GRK3 R587Q as the template and 5’TTCCCCAACCAGCTCGAGTGGC as the forward primer and 5’GCCACTCGAGCTGGTTGGGGAA as the reverse primer. GRK2/3-CAAX constructs were generated by inserting the CAAX overhangs using Gibson assembly to GRK2/3 WT, D110A, R587Q and D110A/R587Q. The generated constructs were validated by sequencing at Eurofins Genomics GmbH. The pcDNA3 backbone was used as a control and is referred to as the empty vector (EV).

### Cell culture

CRISPR/Cas9-generated HEK293 knockout cells of GRK2/3/5/6 (ΔQ-GRK), the family knockouts (ΔGRK2/3, ΔGRK5/6) and our CRISPR/Cas9 HEK293 control cells with unaltered GRK expression^14^ were cultured at 37 °C with 5% CO_2_ in Dulbecco’s modified Eagle’s medium (DMEM; Sigma-Aldrich, D6429), complemented with 10% fetal calf serum (Sigma-Aldrich, F7524) and 1% penicillin and streptomycin mixture (Sigma-Aldrich P0781). Cells were passaged every 3-4 days and regularly checked for infections with mycoplasma using the LONZA MycoAlert mycoplasma detection kit (LT07-318).

### Bioluminescence resonance energy transfer (BRET) measurements

The intermolecular BRET measurements were conducted as described before^14^. In short, cells were seeded into 6 cm dishes (ΔQ-GRK, ΔGRK5/6 1.6 × 10^6^ cells; ΔGRK2/3 1.2 × 10^6^ cells) and transfected the following day with 0.5 μg of the indicated GPCR-NLuc, 1 μg of Halo-tag-β-arrestin2 and 0.25 μg of one GRK construct or empty vector (EV), according to the Effectene transfection reagent manual (Qiagen, #301427). The following day, 40,000 cells per well were seeded into poly-D-lysine-coated 96-well plates (Brand, 781965) and the Halo-ligand was added (1:2,000; Promega, G980A). For each transfection, technical replicates were seeded as triplicates and a mock labelling condition was included without the Halo-ligand fluorophore. Before measuring the next day, the cells were washed twice using measuring buffer (140 mM NaCl, 10 mM HEPES, 5.4 mM KCl, 2 mM CaCl2, 1 mM MgCl2; pH 7.3). After aspiration, the NLuc-substrate furimazine (Promega, N157B) in measuring buffer (1:3,500) was added. The measurements were performed with a Synergy Neo2 plate reader (Biotek), the Gen5 software (version 2.09) and a customized filter cube (fitted with a 555 nm dichroic mirror and a 620/15 bandpass filter). First, basal values were measured for 3 min, followed by addition of the indicated agonist and measurement of the stimulated values for 5 min. The human b2AR was stimulated with isoprenaline (Iso; Sigma-Aldrich, I5627, dissolved in water). The human M2R and M5R were stimulated with Acetylcholine (ACh; Sigma-Aldrich, A6625, dissolved in measuring buffer).

In case of the bARK-CT-mediated inhibition of the GRK2-Gβγ interaction^27,30^ and the measured effects on β-arrestin2 recruitment, 1.2 × 10^6^ CRISPR/Cas9 HEK293 control cells were seeded into 6 cm dishes. The cells were transfected as described above with 0.5 μg of M5R-NLuc and 1 μg of Halo-tag-β-arrestin2, in addition to 0.5 μg or 1 μg of bARK-CT or 1 μg EV as a control. The total amount of transfected DNA was adjusted to 2.5 μg with EV, when necessary.

### Analysis and statistics

The measured BRET ratios were labelling corrected by subtraction of the respective mock-labelled condition and subsequently the averaged stimulated values were divided by the respective averaged baseline values. To get the final dynamic Δ net BRET change, the ligand-dependent labelling-corrected BRET change was divided by the vehicle control and calculated as percent changes. These corrected BRET changes were normalized to the maximum value of GRK2- or GRK-CAAX-mediated recruitment, as indicated in the respective figure legends. In the bARK-CT experiment, the Δ net BRET changes were normalized to the β-arrestin2 recruitment at the highest ligand concentration in absence of bARK-CT (EV-transfected condition). All data are shown as mean of at least three independent experiments ± SEM as indicated. Statistical comparisons were made in GraphPad Prism 7.03 using one-way ANOVA and subsequent Turkey’s test. The supplementary bar graphs of the Halo labelling- and vehicle-corrected mean Δ net BRET changes + SEM before (basal) and after stimulation with the indicated ligand were normalized to the basal BRET ratio derived from the EV-transfected condition (Δ net BRET fold change). Here, statistical differences within one condition between basal and stimulated or between differently transfected conditions were tested using two-way ANOVA followed by a Sidak’s or Tukey’s test respectively. In all cases, a type I error probability of 0.05 was considered significant.

## Data availability

All data supporting the findings of this work are available within the article and supplementary information files.

## Acknowledgements

We want to thank Professor Silvio Gutkind for kindly providing the bARK-CT plasmid.

## Funding

This work was supported by the European Regional Development Fund (Grant ID: EFRE HSB 2018 0019) and by ONCORNET2.0 - H2020-MSCA-ITN-2019; Grant Agreement number: 860229 to C.H. A.G. and J.D. were supported by the University Hospital Jena IZKF (Grant ID: MSP 11 and MSP 10). N.J. was funded with a PhD fellowship from the IZKF, University Hospital Jena. J.C.F. was supported by the Jena School of Molecular Medicine funded by the Carl-Zeiss-Stiftung.

## Author contributions

C.H. developed the concept of the study. E.S.F.M., J.C.F., N.J. performed and analyzed all measurements. J.D. engineered the GRK knockout cells. E.S.F.M., M.R., A.G., N.Y., J.C.F. and C.H. wrote the manuscript with contributions and critical feedback from all authors.

## Competing interests

The authors declare no competing interests.

## Notes

### Competing Interest Statement

The authors have declared no competing interest.

