## Supplementary data file for "GRK specificity and Gβγ dependency determines a GPCR’s potential in biased agonism"

### Supplementary information

**This .pdf includes:**

Supplementary Figure 1-4

Supplementary Table 1-11

**Supplementary Figure 1**

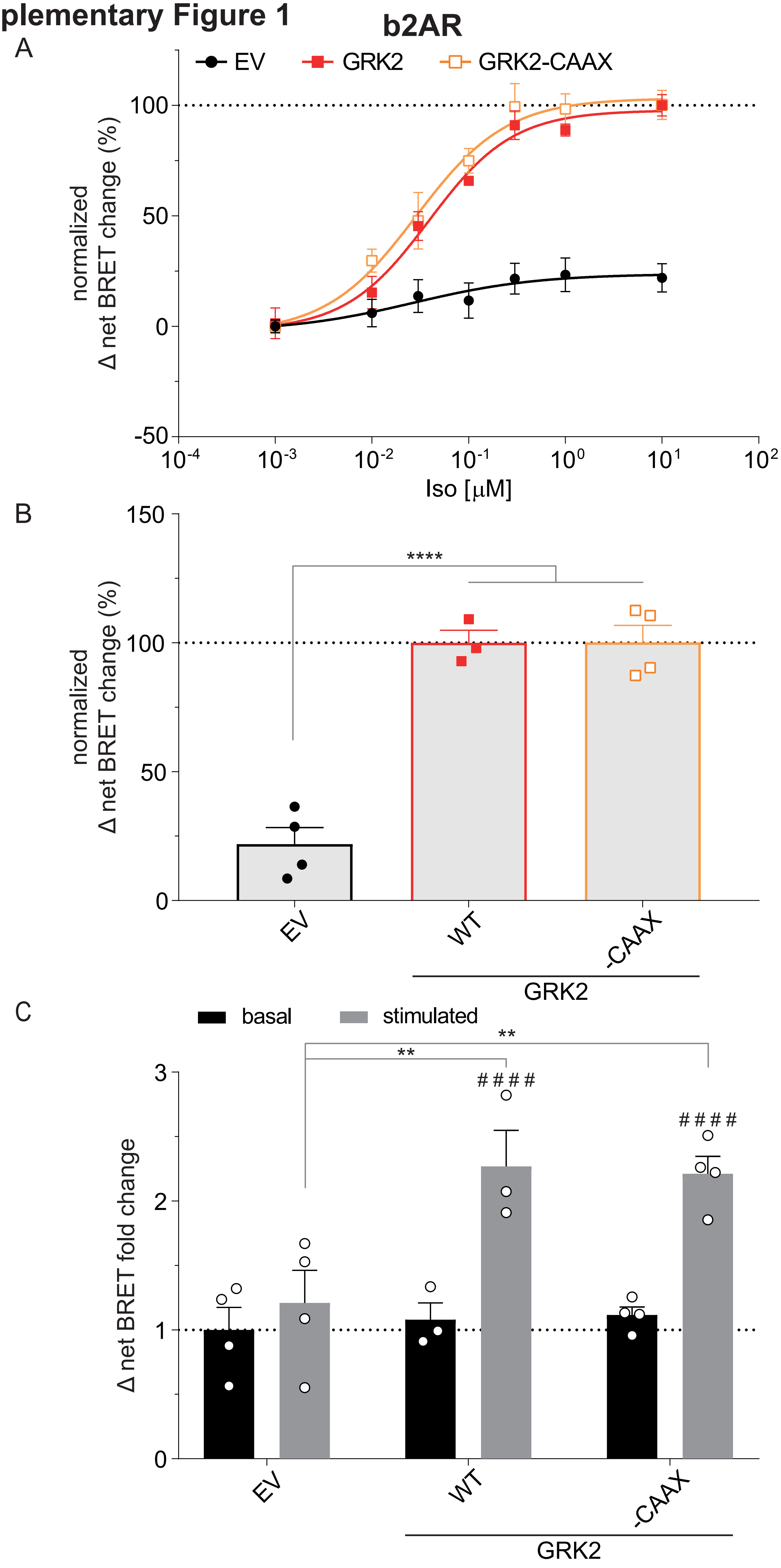

**Suppl. Fig.1: Comparison of GRK2- and GRK2-CAAX-mediated β-arrestin2 recruitment to the b2AR. A,** Iso-induced Halo-Tag-β-arrestin2 recruitment to b2AR-NLuc in ΔQ-GRK cells in absence of the ubiquitously expressed GRKs (empty vector (EV)-transfected) and in presence of wild type GRK2 or GRK2-CAAX. Data from **Fig. 1** are shown again to allow a direct comparison of the GRK2 and GRK2-CAAX condition. Data are shown as Δ net BRET change (%) of at least $n$=3 independent experiments ± SEM, normalized to the maximum response with GRK2. **B,** Normalized BRET data of the highest stimulation of A (10 µM Iso) are displayed as bar graphs. Statistical differences were tested using one-way ANOVA, followed by a Tukey's test (**** $p < 0.0001$). **C**, The Halo-corrected mean Δ net BRET fold changes + SEM of the same dataset (not normalized to the highest stimulated value of the GRK2 WT condition) are presented as bar graphs before (basal) and after stimulation with 10 µM Iso (stimulated). The data were normalized to the basal BRET ratio derived from the EV-transfected condition (dotted line). Statistical differences within one condition between basal and stimulated (#) or between the differently transfected conditions (*) were tested using two-way ANOVA, followed by a Sidak's or Tukey's test respectively (** $p < 0.01$; *** $p < 0.001$; ####/**** $p < 0.0001$). All detailed statistical results are provided in Suppl. Tab. 2.

### Supplementary Figure 2

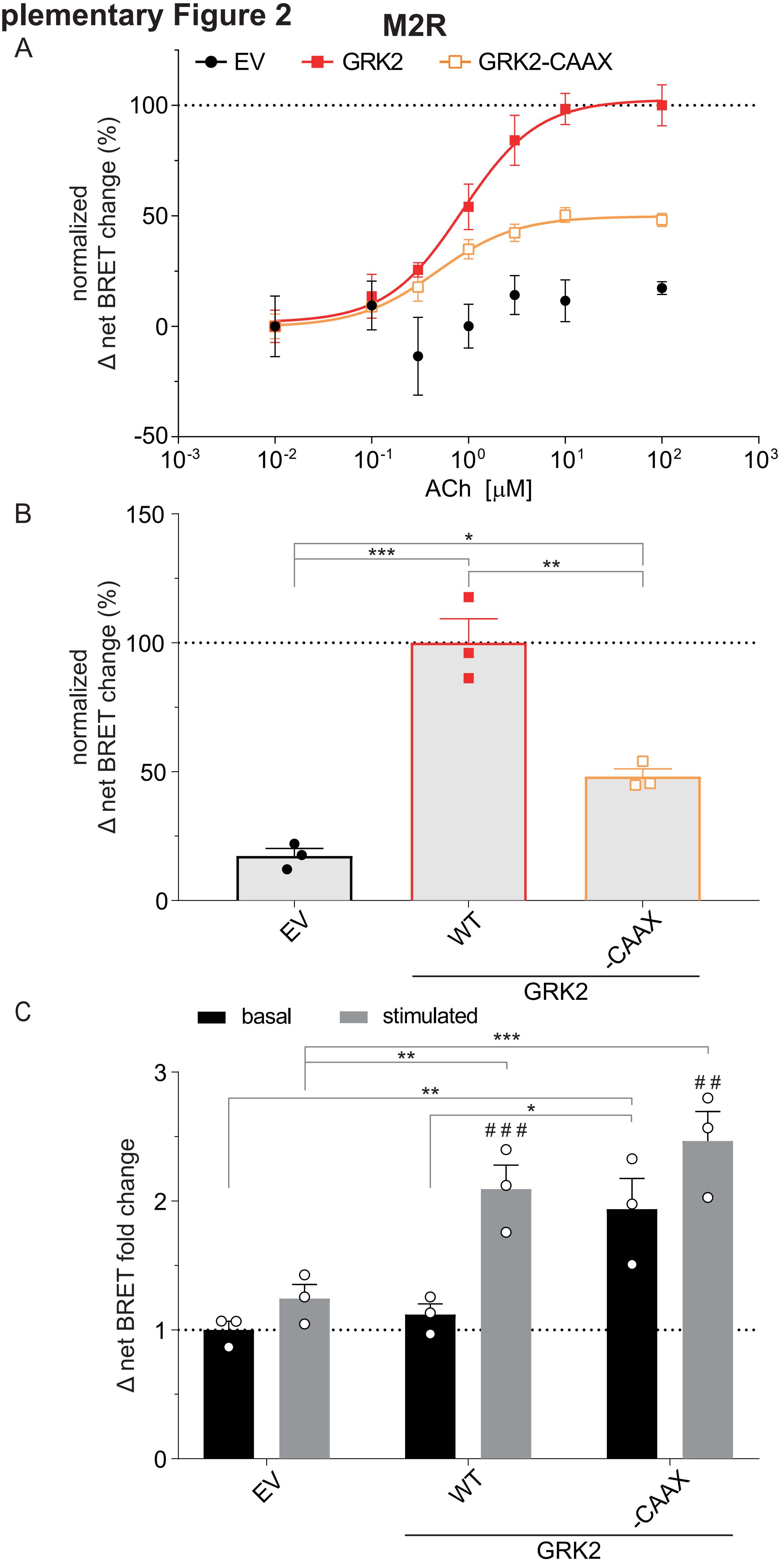

**Suppl. Fig.2: Comparison of GRK2- and GRK2-CAAX-mediated β-arrestin2 recruitment to the M2R. A,** ACh-induced Halo-Tag-β-arrestin2 recruitment to M2R-NLuc in ΔQ-GRK cells in absence of the ubiquitously expressed GRKs (empty vector (EV)-transfected) and in presence of wild type GRK2 or GRK2-CAAX. The concentration-response curves of the data in **Fig. 2 A, B** are shown to allow a direct comparison of the GRK2 and GRK2-CAAX condition. Data are shown as Δ net BRET change (%) of *n*=3 independent experiments ± SEM, normalized to the maximum response with GRK2. **B,** Normalized BRET data of the highest stimulation of A (100 µM ACh) are displayed as bar graphs and statistical differences were tested using one-way ANOVA, followed by a Tukey's test (* $p < 0.05$; ** $p < 0.01$; *** $p < 0.001$). **C**, The Halo-corrected mean Δ net BRET fold changes + SEM of the same dataset (not normalized to the highest stimulated value of the GRK2 WT condition) are presented as bar graphs before (basal) and after stimulation with 100 µM ACh (stimulated). The data were normalized to the basal BRET ratio derived from the EV-transfected condition (dotted line). Statistical differences within one condition between basal and stimulated (#) or between the differently transfected conditions (*) were tested using two-way ANOVA, followed by a Sidak's or Tukey's test respectively (* $p < 0.05$; ##/** $p < 0.01$; ###/*** $p < 0.001$; **** $p < 0.0001$). All detailed statistical results are provided in Suppl. Tab. 5.

**Supplementary Figure 3**

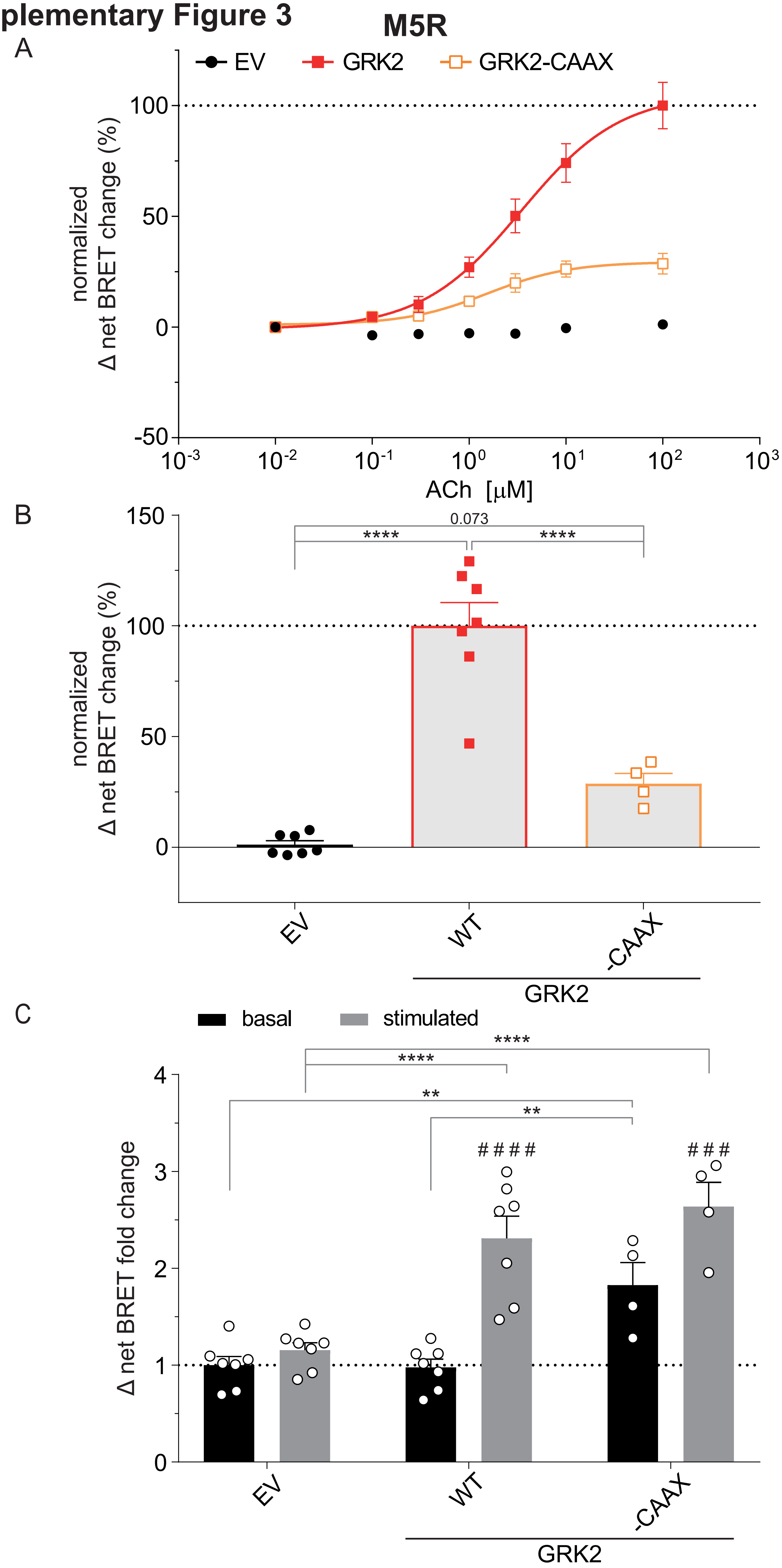

**Suppl. Fig.3: Comparison of GRK2- and GRK2-CAAX-mediated β-arrestin2 recruitment to the M5R. A,** ACh-induced Halo-Tag-β-arrestin2 recruitment to M5R-NLuc in ΔQ-GRK cells in absence of the ubiquitously expressed GRKs (empty vector (EV)-transfected) and in presence of wild type GRK2 or GRK2-CAAX. The concentration-response curves of the data in **Fig. 2 C, D** are shown to allow a direct comparison of the GRK2 and GRK2-CAAX condition. Data are shown as Δ net BRET change (%) of $n$=3 independent experiments ± SEM, normalized to the maximum response with GRK2. **B,** Normalized BRET data of the highest stimulation of A (100 µM ACh) are displayed as bar graphs and statistical differences were tested using one-way ANOVA, followed by a Tukey's test (**** $p < 0.0001$). **C**, The Halo-corrected mean Δ net BRET fold changes + SEM of the same dataset (not normalized to the highest stimulated value of the GRK2 WT condition) are presented as bar graphs before (basal) and after stimulation with 100 µM ACh (stimulated). The data were normalized to the basal BRET ratio derived from the EV-transfected condition (dotted line). Statistical differences within one condition between basal and stimulated (#) or between the differently transfected conditions (*) were tested using two-way ANOVA, followed by a Sidak's or Tukey's test respectively (** $p < 0.01$; ###/*** $p < 0.001$; ####/**** $p < 0.0001$). All detailed statistical results are provided in Suppl. Tab. 6.

### Supplementary Figure 4

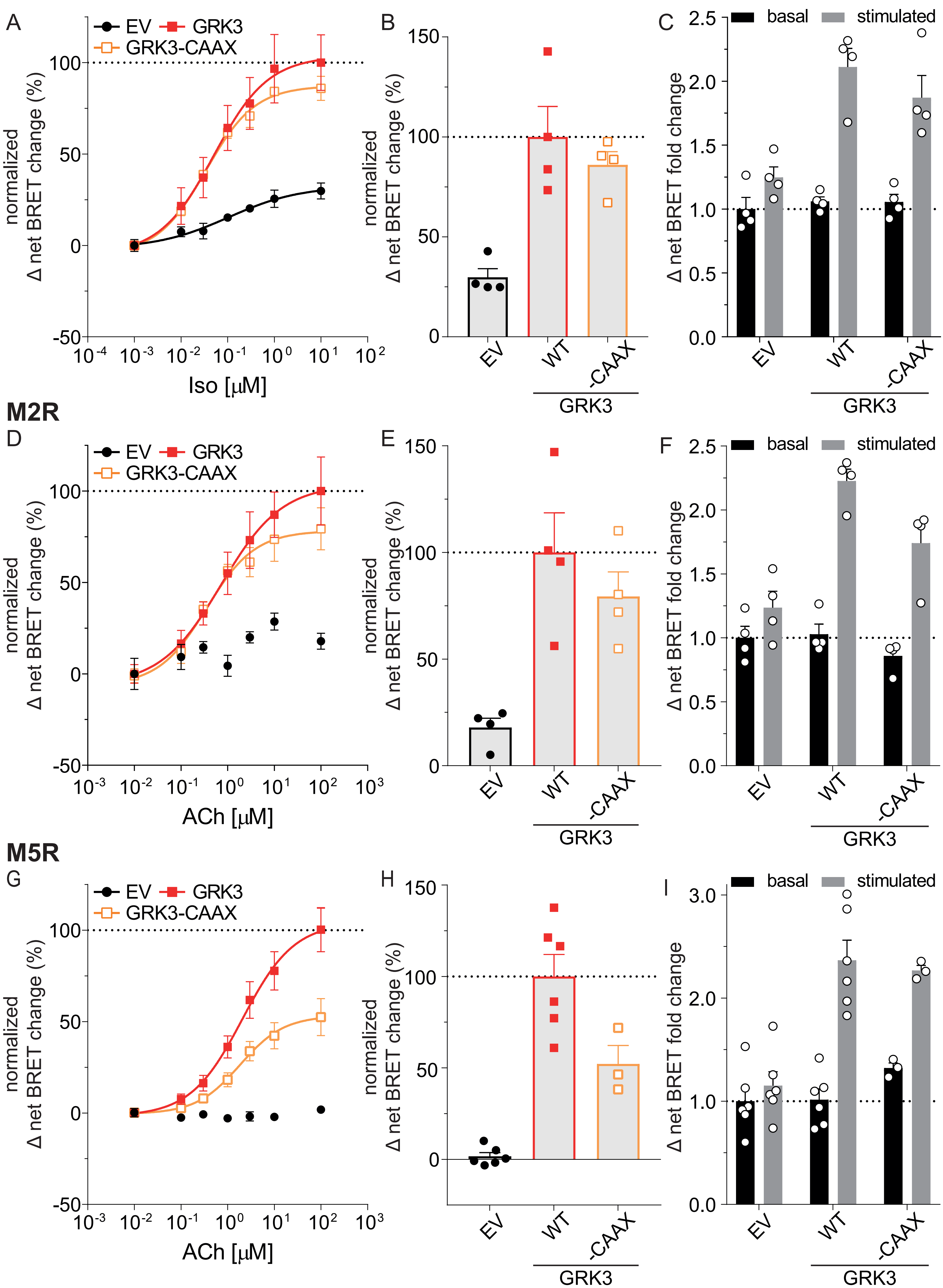

**Suppl. Fig.4: Comparison of GRK3- and GRK3-CAAX-mediated β-arrestin2 recruitment to the b2AR, M2R and M5R. A, D, G,** Agonist-induced Halo-Tag-β-arrestin2 recruitment to b2AR-, M2R- or M5R-NLuc in ΔQ-GRK cells in absence of the ubiquitously expressed GRKs (empty vector (EV)-transfected) and in presence of wild type GRK3 or GRK3-CAAX. The concentration-response curves of the data in **Fig. 3** are shown to allow a direct comparison of the GRK3 and GRK3-CAAX condition. Data are shown as Δ net BRET change (%) of at least $n=3$ independent experiments ± SEM, normalized to the maximum response with GRK3. **B, E, H,** Normalized BRET data of the highest stimulation of A, D, G are displayed as bar graphs and statistical differences were tested using one-way ANOVA, followed by a Tukey's test. **C, F, I,** The Halo-corrected mean Δ net BRET fold changes + SEM of the same dataset (not normalized to the highest stimulated value of the GRK2 WT condition) are presented as bar graphs before (basal) and after stimulation with 10 µM Iso or 100 µM ACh (stimulated). The data were normalized to the basal BRET ratio derived from the EV-transfected condition (dotted line). Statistical differences within one condition between basal and stimulated or between the differently transfected conditions were tested using two-way ANOVA, followed by a Sidak's or Tukey's test respectively. All detailed statistical results are provided in Suppl. Tab. 10.

**Supplementary Table 1:** Detailed statistical results of the analysis of the Δ net BRET change in β-arrestin2 recruitment to b2AR, mediated by the indicated GRK construct in ΔQ-GRK cells, are shown as presented in Figure 1 H and I. The Δ net BRET changes at 10 µM Isoprenaline (Iso) of each condition were compared using one-way ANOVA, followed by a Tukey's test (ns not significant; * p < 0.05; ** p < 0.01; *** p < 0.001; **** p < 0.0001). For each condition the mean difference, 95% confidence interval (CI) of the difference, and the adjusted p value are shown.

| conditions | mean diff. | 95% CI of diff. | adjusted *p* value |
|---|---|---|---|
| EV vs. GRK2 | -78.11 | -106 to -50.18 | <0.0001 **** |
| EV vs. GRK2-D110A | -82.49 | -110.4 to -54.57 | <0.0001 **** |
| EV vs. GRK2-R587Q | -9.005 | -34.86 to 16.85 | 0.8052 ns |
| EV vs. GRK2-D110A,R587Q | -9.693 | -35.55 to 16.16 | 0.762 ns |
| GRK2 vs. GRK2-D110A | -4.386 | -34.24 to 25.47 | 0.9895 ns |
| GRK2 vs. GRK2-R587Q | 69.1 | 41.18 to 97.03 | <0.0001 **** |
| GRK2 vs. GRK2-D110A,R587Q | 68.41 | 40.49 to 96.34 | <0.0001 **** |
| GRK2-D110A vs. GRK2-R587Q | 73.49 | 45.56 to 101.4 | <0.0001 **** |
| GRK2-D110A vs. GRK2-D110A,R587Q | 72.8 | 44.87 to 100.7 | <0.0001 **** |
| GRK2-R587Q vs. GRK2-D110A,R587Q | -0.6872 | -26.54 to 25.17 | >0.9999 ns |
| EV vs. GRK2-CAAX | -78.15 | -108.8 to -47.54 | <0.0001 **** |
| EV vs. GRK2-D110A-CAAX | -75.07 | -105.7 to -44.46 | <0.0001 **** |
| EV vs. GRK2-R587Q-CAAX | -60.52 | -91.13 to -29.92 | 0.0002 *** |
| EV vs. GRK2-D110A,R587Q-CAAX | -62.7 | -93.31 to -32.1 | 0.0001 *** |
| GRK2-CAAX vs. GRK2-D110A-CAAX | 3.08 | -27.53 to 33.69 | 0.9977 ns |
| GRK2-CAAX vs. GRK2-R587Q-CAAX | 17.62 | -12.98 to 48.23 | 0.4204 ns |
| GRK2-CAAX vs. GRK2-D110A,R587Q-CAAX | 15.45 | -15.16 to 46.05 | 0.5434 ns |
| GRK2-D110A-CAAX vs. GRK2-R587Q-CAAX | 14.54 | -16.06 to 45.15 | 0.597 ns |
| GRK2-D110A-CAAX vs. GRK2-D110A,R587Q-CAAX | 12.37 | -18.24 to 42.97 | 0.7249 ns |
| GRK2-R587Q-CAAX vs. GRK2-D110A,R587Q-CAAX | -2.177 | -32.78 to 28.43 | 0.9994 ns |

**Supplementary Table 2:** Detailed statistical results of the analysis of the Δ net BRET change in β-arrestin2 recruitment to b2AR, mediated by the indicated GRK2 construct in ΔQ-GRK cells, are shown as presented in Supplementary Figure 1 B and C. The Δ net BRET changes at 10 µM Isoprenaline (Iso) of each condition were compared using one-way ANOVA, followed by a Tukey's test (ns not significant; * $p < 0.05$; ** $p < 0.01$; *** $p < 0.001$; **** $p < 0.0001$). The non-normalized, Halo-corrected mean Δ net BRET fold changes before (basal) and after stimulation with 10 µM Iso (stimulated) were compared between the conditions or within one condition, as indicated. Statistical analysis was performed by two-way ANOVA, followed by a Tukey's or Sidak's test respectively (ns not significant; * $p < 0.05$; ** $p < 0.01$; *** $p < 0.001$; **** $p < 0.0001$). For each condition the mean difference, 95% confidence interval (CI) of the difference, and the adjusted p value are shown.

Statistical details of Suppl. Fig. 1B

| conditions | mean diff. | 95% CI of diff. | adjusted *p* value | |
|---|---|---|---|---|
| EV vs. GRK2 | -78.11 | -104.4 to -51.81 | <0.0001 | **** |
| EV vs. GRK2 CAAX | -78.3 | -102.6 to -53.95 | <0.0001 | **** |
| GRK2 vs. GRK2 CAAX | -0.1926 | -26.49 to 26.11 | 0.9998 | ns |

Statistical details of Suppl. Fig. 1C

comparison of basal and stimulated between conditions

| basal | mean diff. | 95% CI of diff. | adjusted *p* value | |
|---|---|---|---|---|
| EV vs. GRK2 | -0.07946 | -0.7695 to 0.6106 | 0.9527 | ns |
| EV vs. GRK2 CAAX | -0.1161 | -0.755 to 0.5227 | 0.8866 | ns |
| GRK2 vs. GRK2 CAAX | -0.03669 | -0.7268 to 0.6534 | 0.9897 | ns |

| stimulated | mean diff. | 95% CI of diff. | adjusted *p* value | |
|---|---|---|---|---|
| EV vs. GRK2 | -1.058 | -1.749 to -0.3684 | 0.0031 | ** |
| EV vs. GRK2 CAAX | -1.002 | -1.641 to -0.3627 | 0.0026 | ** |
| GRK2 vs. GRK2 CAAX | 0.05685 | -0.6332 to 0.7469 | 0.9754 | ns |

comparison of basal vs. stimulated within one condition

| conditions | mean diff. | 95% CI of diff. | adjusted *p* value | |
|---|---|---|---|---|
| EV | -0.2095 | -0.5687 to 0.1498 | 0.3136 | ns |
| GRK2 | -1.188 | -1.603 to -0.7737 | <0.0001 | **** |
| GRK2 CAAX | -1.095 | -1.454 to -0.7357 | <0.0001 | **** |

**Supplementary Table 3:** Detailed statistical results of the analysis of the Δ net BRET change in β-arrestin2 recruitment to M2R, mediated by the indicated GRK construct in ΔQ-GRK cells, are shown as presented in Figure 2 A and B. The Δ net BRET changes at 100 µM Acetylcholine (ACh) of each condition were compared using one-way ANOVA, followed by a Tukey's test (ns not significant; * $p < 0.05$; ** $p < 0.01$; *** $p < 0.001$). For each condition the mean difference, 95% confidence interval (CI) of the difference, and the adjusted p value are shown.

| conditions | mean diff. | 95% CI of diff. | adjusted *p* value | |
|---|---|---|---|---|
| EV vs. GRK2 | -82.71 | -116 to -49.41 | <0.0001 | **** |
| EV vs. GRK2-D110A | -103.9 | -137.2 to -70.63 | <0.0001 | **** |
| EV vs. GRK2-R587Q | -6.587 | -39.89 to 26.72 | 0.9626 | ns |
| EV vs. GRK2-D110A,R587Q | -31.28 | -64.59 to 2.019 | 0.0681 | ns |
| GRK2 vs. GRK2-D110A | -21.22 | -54.52 to 12.09 | 0.2924 | ns |
| GRK2 vs. GRK2-R587Q | 76.13 | 42.82 to 109.4 | 0.0002 | *** |
| GRK2 vs. GRK2-D110A,R587Q | 51.43 | 18.13 to 84.73 | 0.0034 | ** |
| GRK2-D110A vs. GRK2-R587Q | 97.34 | 64.04 to 130.6 | <0.0001 | **** |
| GRK2-D110A vs. GRK2-D110A,R587Q | 72.65 | 39.34 to 105.9 | 0.0002 | *** |
| GRK2-R587Q vs. GRK2-D110A,R587Q | -24.7 | -58 to 8.606 | 0.1814 | ns |
| EV vs. GRK2-CAAX | -64.07 | -116.7 to -11.43 | 0.0166 | * |
| EV vs. GRK2-D110A-CAAX | -63.34 | -116 to -10.7 | 0.0178 | * |
| EV vs. GRK2-R587Q-CAAX | -59.71 | -112.4 to -7.066 | 0.0252 | * |
| EV vs. GRK2-D110A,R587Q-CAAX | -113.8 | -166.4 to -61.16 | 0.0002 | *** |
| GRK2-CAAX vs. GRK2-D110A-CAAX | 0.7327 | -51.91 to 53.37 | >0.9999 | ns |
| GRK2-CAAX vs. GRK2-R587Q-CAAX | 4.362 | -48.28 to 57 | 0.9986 | ns |
| GRK2-CAAX vs. GRK2-D110A,R587Q-CAAX | -49.73 | -102.4 to 2.915 | 0.0663 | ns |
| GRK2-D110A-CAAX vs. GRK2-R587Q-CAAX | 3.629 | -49.01 to 56.27 | 0.9993 | ns |
| GRK2-D110A-CAAX vs. GRK2-D110A,R587Q-CAAX | -50.46 | -103.1 to 2.183 | 0.0618 | ns |
| GRK2-R587Q-CAAX vs. GRK2-D110A,R587Q-CAAX | -54.09 | -106.7 to -1.447 | 0.0434 | * |

**Supplementary Table 4:** Detailed statistical results of the analysis of the Δ net BRET change in β-arrestin2 recruitment to M5R, mediated by the indicated GRK2 construct in ΔQ-GRK cells, are shown as presented in Figure 2 C and D. The Δ net BRET changes at 100 µM ACh of each condition were compared using one-way ANOVA, followed by a Tukey's test (ns not significant; * $p < 0.05$; ** $p < 0.01$; *** $p < 0.001$; **** $p < 0.0001$). For each condition the mean difference, 95% confidence interval (CI) of the difference, and the adjusted p value are shown.

| conditions | mean diff. | 95% CI of diff. | adjusted *p* value | |
|---|---|---|---|---|
| EV vs. GRK2 | -98.85 | -128.9 to -68.81 | <0.0001 | **** |
| EV vs. GRK2-D110A | -87.82 | -119.1 to -56.56 | <0.0001 | **** |
| EV vs. GRK2-R587Q | -41.19 | -72.45 to -9.923 | 0.0055 | ** |
| EV vs. GRK2-D110A,R587Q | -19.88 | -51.14 to 11.39 | 0.3637 | ns |
| GRK2 vs. GRK2-D110A | 11.03 | -20.24 to 42.29 | 0.8392 | ns |
| GRK2 vs. GRK2-R587Q | 57.66 | 26.4 to 88.93 | <0.0001 | **** |
| GRK2 vs. GRK2-D110A,R587Q | 78.97 | 47.71 to 110.2 | <0.0001 | **** |
| GRK2-D110A vs. GRK2-R587Q | 46.64 | 14.19 to 79.08 | 0.0022 | ** |
| GRK2-D110A vs. GRK2-D110A,R587Q | 67.95 | 35.5 to 100.4 | <0.0001 | **** |
| GRK2-R587Q vs. GRK2-D110A,R587Q | 21.31 | -11.13 to 53.75 | 0.3322 | ns |
| EV vs. GRK2-CAAX | -95.98 | -165.3 to -26.7 | 0.0046 | ** |
| EV vs. GRK2-D110A-CAAX | -112.7 | -189 to -36.41 | 0.0026 | ** |
| EV vs. GRK2-R587Q-CAAX | -107.4 | -176.6 to -38.07 | 0.0016 | ** |
| EV vs. GRK2-D110A,R587Q-CAAX | -113.3 | -182.6 to -44.05 | 0.001 | *** |
| GRK2-CAAX vs. GRK2-D110A-CAAX | -16.71 | -101.1 to 67.72 | 0.9728 | ns |
| GRK2-CAAX vs. GRK2-R587Q-CAAX | -11.37 | -89.53 to 66.79 | 0.9913 | ns |
| GRK2-CAAX vs. GRK2-D110A,R587Q-CAAX | -17.36 | -95.52 to 60.81 | 0.9591 | ns |
| GRK2-D110A-CAAX vs. GRK2-R587Q-CAAX | 5.336 | -79.09 to 89.76 | 0.9997 | ns |
| GRK2-D110A-CAAX vs. GRK2-D110A,R587Q-CAAX | -0.6486 | -85.07 to 83.77 | >0.9999 | ns |
| GRK2-R587Q-CAAX vs. GRK2-D110A,R587Q-CAAX | -5.985 | -84.15 to 72.18 | 0.9993 | ns |

**Supplementary Table 5:** Detailed statistical results of the analysis of the Δ net BRET change in β-arrestin2 recruitment to M2R, mediated by the indicated GRK2 construct in ΔQ-GRK cells, are shown as presented in Supplementary Figure 2 B and C. The Δ net BRET changes at 100 µM ACh of each condition were compared using one-way ANOVA, followed by a Tukey's test (ns not significant; * $p < 0.05$; ** $p < 0.01$; *** $p < 0.001$; **** $p < 0.0001$). The non-normalized, Halo-corrected mean Δ net BRET fold changes before (basal) and after stimulation with 100 µM ACh (stimulated) were compared between the conditions or within one condition, as indicated. Statistical analysis was performed by two-way ANOVA, followed by a Tukey's or Sidak's test respectively (ns not significant; * $p < 0.05$; ** $p < 0.01$; *** $p < 0.001$; **** $p < 0.0001$). For each condition the mean difference, 95% confidence interval (CI) of the difference, and the adjusted p value are shown.

Statistical details of Suppl. Fig. 2B

| conditions | mean diff. | 95% CI of diff. | adjusted *p* value | |
|---|---|---|---|---|
| EV vs. GRK2 | -82.71 | -108.2 to -57.24 | 0.0001 | *** |
| EV vs. GRK2 CAAX | -30.83 | -56.3 to -5.353 | 0.0232 | * |
| GRK2 vs. GRK2 CAAX | 51.89 | 26.42 to 77.36 | 0.0019 | ** |

Statistical details of Suppl. Fig. 2C

comparison of basal and stimulated between conditions

| basal | mean diff. | 95% CI of diff. | adjusted *p* value | |
|---|---|---|---|---|
| EV vs. GRK2 | -0.1183 | -0.7474 to 0.5108 | 0.8719 | ns |
| EV vs. GRK2 CAAX | -0.9371 | -1.566 to -0.308 | 0.0049 | ** |
| GRK2 vs. GRK2 CAAX | -0.8188 | -1.448 to -0.1897 | 0.0119 | * |

| stimulated | mean diff. | 95% CI of diff. | adjusted *p* value | |
|---|---|---|---|---|
| EV vs. GRK2 | -0.8504 | -1.48 to -0.2213 | 0.0093 | ** |
| EV vs. GRK2 CAAX | -1.223 | -1.852 to -0.5937 | 0.0006 | *** |
| GRK2 vs. GRK2 CAAX | -0.3724 | -1.001 to 0.2567 | 0.2915 | ns |

comparison of basal vs. stimulated within one condition

| conditions | mean diff. | 95% CI of diff. | adjusted *p* value | |
|---|---|---|---|---|
| EV | -0.2422 | -0.551 to 0.06652 | 0.1219 | ns |
| GRK2 | -0.9743 | -1.283 to -0.6656 | 0.0001 | *** |
| GRK2 CAAX | -0.528 | -0.8367 to -0.2192 | 0.0041 | ** |

**Supplementary Table 6:** Detailed statistical results of the analysis of the Δ net BRET change in β-arrestin2 recruitment to M5R, mediated by the indicated GRK2 construct in ΔQ-GRK cells, are shown as presented in Supplementary Figure 3 B and C. The Δ net BRET changes at 100 µM ACh of each condition were compared using one-way ANOVA, followed by a Tukey's test (ns not significant; * $p < 0.05$; ** $p < 0.01$; *** $p < 0.001$; **** $p < 0.0001$). The non-normalized, Halo-corrected mean Δ net BRET fold changes before (basal) and after stimulation with 100 µM ACh (stimulated) were compared between the conditions or within one condition, as indicated. Statistical analysis was performed by two-way ANOVA, followed by a Tukey's or Sidak's test respectively (ns not significant; * $p < 0.05$; ** $p < 0.01$; *** $p < 0.001$; **** $p < 0.0001$). For each condition the mean difference, 95% confidence interval (CI) of the difference, and the adjusted p value are shown.

Statistical details of Suppl. Fig. 3B

| conditions | mean diff. | 95% CI of diff. | adjusted *p* value | |
|---|---|---|---|---|
| EV vs. GRK2 | -98.85 | -124.3 to -73.39 | <0.0001 | **** |
| EV vs. GRK2 CAAX | -27.49 | -57.34 to 2.361 | 0.0733 | ns |
| GRK2 vs. GRK2 CAAX | 71.36 | 41.51 to 101.2 | <0.0001 | **** |

Statistical details of Suppl. Fig. 3C

comparison of basal and stimulated between conditions

| basal | mean diff. | 95% CI of diff. | adjusted *p* value | |
|---|---|---|---|---|
| EV vs. GRK2 | 0.02301 | -0.4865 to 0.5325 | 0.9932 | ns |
| EV vs. GRK2 CAAX | -0.8273 | -1.425 to -0.2298 | 0.0051 | ** |
| GRK2 vs. GRK2 CAAX | -0.8503 | -1.448 to -0.2528 | 0.004 | ** |

| stimulated | mean diff. | 95% CI of diff. | adjusted *p* value | |
|---|---|---|---|---|
| EV vs. GRK2 | -1.152 | -1.662 to -0.6429 | <0.0001 | **** |
| EV vs. GRK2 CAAX | -1.481 | -2.079 to -0.8839 | <0.0001 | **** |
| GRK2 vs. GRK2 CAAX | -0.329 | -0.9264 to 0.2685 | 0.3755 | ns |

comparison of basal vs. stimulated within one condition

| conditions | mean diff. | 95% CI of diff. | adjusted *p* value | |
|---|---|---|---|---|
| EV | -0.1559 | -0.4621 to 0.1503 | 0.472 | ns |
| GRK2 | -1.331 | -1.638 to -1.025 | <0.0001 | **** |
| GRK2 CAAX | -0.81 | -1.215 to -0.4049 | 0.0002 | *** |

**Supplementary Table 7:** Detailed statistical results of the analysis of the Δ net BRET change in β-arrestin2 recruitment to b2AR, mediated by the indicated GRK3 construct in ΔQ-GRK cells, are shown as presented in Figure 3 A and B. The Δ net BRET changes at 10 μM Iso of each condition were compared using one-way ANOVA, followed by a Tukey's test (ns not significant; * $p < 0.05$; ** $p < 0.01$; *** $p < 0.001$; **** $p < 0.0001$). For each condition the mean difference, 95% confidence interval (CI) of the difference, and the adjusted p value are shown.

| conditions | mean diff. | 95% CI of diff. | adjusted *p* value | |
|---|---|---|---|---|
| EV vs. GRK3 | -70.23 | -115.2 to -25.21 | 0.002 | ** |
| EV vs. GRK3-D110A | -64.53 | -113.1 to -15.9 | 0.0075 | ** |
| EV vs. GRK3-R587Q | 0.298 | -44.72 to 45.31 | >0.9999 | ns |
| EV vs. GRK3-D110A,R587Q | 4.095 | -40.92 to 49.11 | 0.9984 | ns |
| GRK3 vs. GRK3-D110A | 5.7 | -42.92 to 54.32 | 0.9957 | ns |
| GRK3 vs. GRK3-R587Q | 70.52 | 25.51 to 115.5 | 0.0019 | ** |
| GRK3 vs. GRK3-D110A,R587Q | 74.32 | 29.31 to 119.3 | 0.0012 | ** |
| GRK3-D110A vs. GRK3-R587Q | 64.82 | 16.2 to 113.4 | 0.0073 | ** |
| GRK3-D110A vs. GRK3-D110A,R587Q | 68.62 | 20 to 117.2 | 0.0046 | ** |
| GRK3-R587Q vs. GRK3-D110A,R587Q | 3.797 | -41.22 to 48.81 | 0.9988 | ns |
| EV vs. GRK3-CAAX | -65.39 | -91.19 to -39.58 | <0.0001 | **** |
| EV vs. GRK3-D110A-CAAX | -50.44 | -76.24 to -24.63 | 0.0002 | *** |
| EV vs. GRK3-R587Q-CAAX | -40.91 | -68.78 to -13.03 | 0.0033 | ** |
| EV vs. GRK3-D110A,R587Q-CAAX | -17.57 | -43.37 to 8.24 | 0.2649 | ns |
| GRK3-CAAX vs. GRK3-D110A-CAAX | 14.95 | -10.86 to 40.75 | 0.4085 | ns |
| GRK3-CAAX vs. GRK3-R587Q-CAAX | 24.48 | -3.393 to 52.35 | 0.098 | ns |
| GRK3-CAAX vs. GRK3-D110A,R587Q-CAAX | 47.82 | 22.02 to 73.63 | 0.0004 | *** |
| GRK3-D110A-CAAX vs. GRK3-R587Q-CAAX | 9.531 | -18.34 to 37.4 | 0.8207 | ns |
| GRK3-D110A-CAAX vs. GRK3-D110A,R587Q-CAAX | 32.87 | 7.067 to 58.68 | 0.0103 | * |
| GRK3-R587Q-CAAX vs. GRK3-D110A,R587Q-CAAX | 23.34 | -4.531 to 51.21 | 0.1219 | ns |

**Supplementary Table 8:** Detailed statistical results of the analysis of the Δ net BRET change in β-arrestin2 recruitment to M2R, mediated by the indicated GRK3 construct in ΔQ-GRK cells, are shown as presented in Figure 3 C and D. The Δ net BRET changes at 100 µM ACh of each condition were compared using one-way ANOVA, followed by a Tukey's test (ns not significant; * $p < 0.05$; ** $p < 0.01$; *** $p < 0.001$; **** $p < 0.0001$). For each condition the mean difference, 95% confidence interval (CI) of the difference, and the adjusted p value are shown.

| conditions | mean diff. | 95% CI of diff. | adjusted *p* value | |
|---|---|---|---|---|
| EV vs. GRK3 | -82.13 | -133.2 to -31.1 | 0.0015 | ** |
| EV vs. GRK3-D110A | -62.4 | -113.4 to -11.37 | 0.0138 | * |
| EV vs. GRK3-R587Q | 11.8 | -39.23 to 62.84 | 0.9482 | ns |
| EV vs. GRK3-D110A,R587Q | -13.03 | -68.15 to 42.09 | 0.9441 | ns |
| GRK3 vs. GRK3-D110A | 19.74 | -31.29 to 70.77 | 0.7487 | ns |
| GRK3 vs. GRK3-R587Q | 93.94 | 42.91 to 145 | 0.0004 | *** |
| GRK3 vs. GRK3-D110A,R587Q | 69.1 | 13.98 to 124.2 | 0.0116 | * |
| GRK3-D110A vs. GRK3-R587Q | 74.2 | 23.17 to 125.2 | 0.0036 | ** |
| GRK3-D110A vs. GRK3-D110A,R587Q | 49.37 | -5.754 to 104.5 | 0.0893 | ns |
| GRK3-R587Q vs. GRK3-D110A,R587Q | -24.84 | -79.95 to 30.28 | 0.6352 | ns |
| EV vs. GRK3-CAAX | -77.48 | -123.4 to -31.53 | 0.0011 | ** |
| EV vs. GRK3-D110A-CAAX | -61.54 | -107.5 to -15.59 | 0.0074 | ** |
| EV vs. GRK3-R587Q-CAAX | -48.05 | -97.68 to 1.586 | 0.0596 | ns |
| EV vs. GRK3-D110A,R587Q-CAAX | -21.72 | -71.35 to 27.92 | 0.6512 | ns |
| GRK3-CAAX vs. GRK3-D110A-CAAX | 15.94 | -30.01 to 61.89 | 0.8074 | ns |
| GRK3-CAAX vs. GRK3-R587Q-CAAX | 29.44 | -20.2 to 79.07 | 0.3799 | ns |
| GRK3-CAAX vs. GRK3-D110A,R587Q-CAAX | 55.77 | 6.133 to 105.4 | 0.025 | * |
| GRK3-D110A-CAAX vs. GRK3-R587Q-CAAX | 13.49 | -36.14 to 63.13 | 0.9078 | ns |
| GRK3-D110A-CAAX vs. GRK3-D110A,R587Q-CAAX | 39.82 | -9.809 to 89.46 | 0.144 | ns |
| GRK3-R587Q-CAAX vs. GRK3-D110A,R587Q-CAAX | 26.33 | -26.73 to 79.39 | 0.5438 | ns |

**Supplementary Table 9:** Detailed statistical results of the analysis of the Δ net BRET change in β-arrestin2 recruitment to M5R, mediated by the indicated GRK3 construct in ΔQ-GRK cells, are shown as presented in Figure 3 E and F . The Δ net BRET changes at 100 µM ACh of each condition were compared using one-way ANOVA, followed by a Tukey's test (ns not significant; * $p < 0.05$; ** $p < 0.01$; *** $p < 0.001$; **** $p < 0.0001$). For each condition the mean difference, 95% confidence interval (CI) of the difference, and the adjusted p value are shown.

| conditions | mean diff. | 95% CI of diff. | adjusted *p* value | |
|---|---|---|---|---|
| EV vs. GRK3 | -98.37 | -130.7 to -66.04 | <0.0001 | **** |
| EV vs. GRK3-D110A | -86.41 | -120.3 to -52.5 | <0.0001 | **** |
| EV vs. GRK3-R587Q | -47.8 | -80.13 to -15.48 | 0.0018 | ** |
| EV vs. GRK3-D110A,R587Q | -9.784 | -42.11 to 22.54 | 0.8972 | ns |
| GRK3 vs. GRK3-D110A | 11.96 | -21.94 to 45.86 | 0.8346 | ns |
| GRK3 vs. GRK3-R587Q | 50.56 | 18.24 to 82.89 | 0.001 | *** |
| GRK3 vs. GRK3-D110A,R587Q | 88.58 | 56.26 to 120.9 | <0.0001 | **** |
| GRK3-D110A vs. GRK3-R587Q | 38.6 | 4.7 to 72.5 | 0.0202 | * |
| GRK3-D110A vs. GRK3-D110A,R587Q | 76.62 | 42.72 to 110.5 | <0.0001 | **** |
| GRK3-R587Q vs. GRK3-D110A,R587Q | 38.02 | 5.695 to 70.34 | 0.0156 | * |
| EV vs. GRK3-CAAX | -96.87 | -158.3 to -35.47 | 0.0018 | ** |
| EV vs. GRK3-D110A-CAAX | -97.22 | -168.1 to -26.31 | 0.0058 | ** |
| EV vs. GRK3-R587Q-CAAX | -142.3 | -198.3 to -86.19 | <0.0001 | **** |
| EV vs. GRK3-D110A,R587Q-CAAX | -83.06 | -139.1 to -27.01 | 0.0031 | ** |
| GRK3-CAAX vs. GRK3-D110A-CAAX | -0.341 | -79.62 to 78.94 | >0.9999 | ns |
| GRK3-CAAX vs. GRK3-R587Q-CAAX | -45.38 | -111.7 to 20.95 | 0.2607 | ns |
| GRK3-CAAX vs. GRK3-D110A,R587Q-CAAX | 13.81 | -52.52 to 80.14 | 0.9641 | ns |
| GRK3-D110A-CAAX vs. GRK3-R587Q-CAAX | -45.04 | -120.2 to 30.17 | 0.3778 | ns |
| GRK3-D110A-CAAX vs. GRK3-D110A,R587Q-CAAX | 14.15 | -61.06 to 89.36 | 0.975 | ns |
| GRK3-R587Q-CAAX vs. GRK3-D110A,R587Q-CAAX | 59.19 | -2.221 to 120.6 | 0.0613 | ns |

**Supplementary Table 10:** Detailed statistical results of the analysis of the Δ net BRET change in β-arrestin2 recruitment to b2AR, M2R and M5R, mediated by the indicated GRK3 construct in ΔQ-GRK cells, are shown as presented in Supplementary Figure 4. The Δ net BRET changes at highest ligand concentration of each condition were compared using one-way ANOVA (Suppl. Fig. 4 B, E and H), followed by a Tukey's test (ns not significant; * p < 0.05; ** p < 0.01; *** p < 0.001; **** p < 0.0001). The non-normalized, Halo-corrected mean Δ net BRET fold changes before (basal) and after stimulation with highest ligand concentration (stimulated) were compared between the conditions or within one condition (Suppl. Fig. 4 C, F and I), as indicated. Statistical analysis was performed by two-way ANOVA, followed by a Tukey's or Sidak's test respectively (ns not significant; * p < 0.05; ** p < 0.01; *** p < 0.001; **** p < 0.0001). For each condition the mean difference, 95% confidence interval (CI) of the difference, and the adjusted p value are shown.

**b2AR**

Statistical details of Suppl. Fig. 4B

| conditions | mean diff. | 95% CI of diff. | adjusted *p* value |
|---|---|---|---|
| EV vs. GRK3 | -70.23 | -109.4 to -31.04 | 0.0019 ** |
| EV vs. GRK3 CAAX | -56.22 | -95.41 to -17.04 | 0.0078 ** |
| GRK3 vs. GRK3 CAAX | 14 | -25.18 to 53.19 | 0.5964 ns |

Statistical details of Suppl. Fig. 4C

comparison of basal and stimulated between conditions

| basal | mean diff. | 95% CI of diff. | adjusted *p* value |
|---|---|---|---|
| EV vs. GRK3 | -0.05886 | -0.4522 to 0.3345 | 0.9231 ns |
| EV vs. GRK3 CAAX | -0.05543 | -0.4488 to 0.3379 | 0.9315 ns |
| GRK3 vs. GRK3 CAAX | 0.003428 | -0.3899 to 0.3968 | 0.9997 ns |

| stimulated | mean diff. | 95% CI of diff. | adjusted *p* value |
|---|---|---|---|
| EV vs. GRK3 | -0.8634 | -1.257 to -0.47 | <0.0001 **** |
| EV vs. GRK3 CAAX | -0.6237 | -1.017 to -0.2303 | 0.0021 ** |
| GRK3 vs. GRK3 CAAX | 0.2397 | -0.1537 to 0.6331 | 0.29 ns |

comparison of basal vs. stimulated within one condition

| conditions | mean diff. | 95% CI of diff. | adjusted *p* value |
|---|---|---|---|
| EV | -0.2476 | -0.561 to 0.06585 | 0.1326 ns |
| GRK3 | -1.052 | -1.366 to -0.7387 | <0.0001 **** |
| GRK3 CAAX | -0.8158 | -1.129 to -0.5024 | <0.0001 **** |

**M2R**

Statistical details of Suppl. Fig. 4E

| conditions | mean diff. | 95% CI of diff. | adjusted *p* value |
|---|---|---|---|
| EV vs. GRK3 | -82.13 | -133.1 to -31.2 | 0.0038 ** |
| EV vs. GRK3 CAAX | -61.49 | -112.4 to -10.56 | 0.0203 * |
| GRK3 vs. GRK3 CAAX | 20.65 | -30.29 to 71.58 | 0.5198 ns |

Statistical details of Suppl. Fig. 4F

comparison of basal and stimulated between conditions

| basal | mean diff. | 95% CI of diff. | adjusted *p* value |
|---|---|---|---|
| EV vs. GRK3 | -0.02757 | -0.4106 to 0.3555 | 0.9816 ns |
| EV vs. GRK3 CAAX | 0.1415 | -0.2416 to 0.5245 | 0.6213 ns |
| GRK3 vs. GRK3 CAAX | 0.169 | -0.214 to 0.5521 | 0.5109 ns |

| stimulated | mean diff. | 95% CI of diff. | adjusted *p* value |
|---|---|---|---|
| EV vs. GRK3 | -0.9904 | -1.373 to -0.6073 | <0.0001 **** |
| EV vs. GRK3 CAAX | -0.5045 | -0.8875 to -0.1214 | 0.0093 ** |
| GRK3 vs. GRK3 CAAX | 0.4859 | 0.1028 to 0.8689 | 0.0121 * |

comparison of basal vs. stimulated within one condition

| conditions | mean diff. | 95% CI of diff. | adjusted *p* value |
|---|---|---|---|
| EV | -0.2365 | -0.455 to -0.01798 | 0.0341 * |
| GRK3 | -1.199 | -1.418 to -0.9808 | <0.0001 **** |
| GRK3 CAAX | -0.8824 | -1.101 to -0.6639 | <0.0001 **** |

**M5R**

Statistical details of Suppl. Fig. 4H

| conditions | mean diff. | 95% CI of diff. | adjusted *p* value |
|---|---|---|---|
| EV vs. GRK3 | -98.37 | -130.2 to -66.55 | <0.0001 **** |
| EV vs. GRK3 CAAX | -50.56 | -89.53 to -11.59 | 0.0121 * |
| GRK3 vs. GRK3 CAAX | 47.81 | 8.843 to 86.77 | 0.017 * |

Statistical details of Suppl. Fig. 4I

comparison of basal and stimulated between conditions

| basal | mean diff. | 95% CI of diff. | adjusted *p* value |
|---|---|---|---|
| EV vs. GRK3 | -0.01621 | -0.4838 to 0.4514 | 0.9959 ns |
| EV vs. GRK3 CAAX | -0.322 | -0.8946 to 0.2507 | 0.3547 ns |
| GRK3 vs. GRK3 CAAX | -0.3058 | -0.8784 to 0.2669 | 0.391 ns |

| stimulated | mean diff. | 95% CI of diff. | adjusted *p* value |
|---|---|---|---|
| EV vs. GRK3 | -1.216 | -1.684 to -0.7485 | <0.0001 **** |
| EV vs. GRK3 CAAX | -1.117 | -1.69 to -0.5444 | 0.0002 *** |
| GRK3 vs. GRK3 CAAX | 0.099 | -0.4737 to 0.6717 | 0.9028 ns |

comparison of basal vs. stimulated within one condition

| conditions | mean diff. | 95% CI of diff. | adjusted *p* value |
|---|---|---|---|
| EV | -0.1512 | -0.3849 to 0.08257 | 0.2672 ns |
| GRK3 | -1.351 | -1.585 to -1.117 | <0.0001 **** |
| GRK3 CAAX | -0.9462 | -1.277 to -0.6157 | <0.0001 **** |

**Supplementary Table 11:** Detailed statistical results of the analysis of the Δ net BRET change in β-arrestin2 recruitment to M5R, mediated by the indicated amount of bARK-CT in CRISPR/Cas9 Control cells, are shown as presented in Figure 4 B. The Δ net BRET changes at 100 µM ACh of each condition were compared using one-way ANOVA, followed by a Tukey's test (ns not significant; * $p < 0.05$; ** $p < 0.01$; *** $p < 0.001$; **** $p < 0.0001$). For each condition the mean difference, 95% confidence interval (CI) of the difference, and the adjusted p value are shown.

| conditions | mean diff. | 95% CI of diff. | adjusted *p* value | |
|---|---|---|---|---|
| EV vs. 0.5 µg bARK-CT | 15.9 | -8.554 to 40.36 | 0.2124 | ns |
| EV vs. 1 µg bARK-CT | 43.84 | 21.23 to 66.45 | 0.0014 | ** |
| 0.5 µg bARK-CT vs. 1 µg bARK-CT | 27.93 | 3.509 to 52.36 | 0.0275 | * |
